# HDAC3 inhibition stabilizes the IL-37 receptor module to enhance anti-inflammatory signaling in cystic fibrosis airway epithelium

**DOI:** 10.64898/2026.06.02.729465

**Authors:** Keiko Ueno-Shuto, Ayami Fukuyama, Kota Nakajima, Yuki Hitora, Sachiko Tsukamoto, Tomoki Kishimoto, Keisuke Kawano, Koji Nishi, Mary Ann Suico, Tsuyoshi Shuto

**Affiliations:** Laboratory of Pharmacology, Division of Life Science, Faculty of Pharmaceutical Sciences, Sojo University, Kumamoto 860-0082, Japan; Department of Molecular Medicine, Graduate School of Pharmaceutical Sciences, Kumamoto University, 5-1 Oe-Honmachi, Chuo-ku, Kumamoto 862-0973, Japan; Department of Natural Medicines, Graduate School of Pharmaceutical Sciences, Kumamoto University, Kumamoto 862-0973, Japan; Program for Fostering Innovators to Lead a Better Co-being Society, Kumamoto University, 2-39-1 Kurokami, Chuo-ku, Kumamoto 862-8555, Japan; Global Center for Natural Resources Sciences, Faculty of Life Sciences, Kumamoto University, 5-1 Oe-Honmachi, Chuo-ku, Kumamoto 862-0973, Japan

**Author notes:** Correspondence and requests for materials should be addressed to T.S.

**Keywords:** HDAC3, SIGIRR, IL-1R8, IL-18Rα, IL-37b, cystic fibrosis, airway epithelium, mucosal immunity, proteostasis

## Abstract

Airway inflammation in cystic fibrosis (CF) persists despite advances in CFTR modulator therapy. IL-37b suppresses innate immune signaling through a receptor complex containing IL-18Rα and wild-type SIGIRR (WT-SIGIRR; IL-1R8), but this pathway is compromised in CF airway epithelial cells by the dominant-negative exon 8-skipped SIGIRR isoform (Δ8-SIGIRR). Here, a natural-product screen identified short-chain fatty acids as preferential enhancers of WT-SIGIRR. Pan-HDAC inhibition with panobinostat increased WT-SIGIRR, reduced Δ8-SIGIRR, and restored IL-37b-dependent suppression of the TLR3 ligand poly(I:C)-induced IL-8 production. Isoform-selective inhibitor screening and siRNA knockdown identified HDAC3 as a regulator of the IL-37 receptor module. Low concentrations of RGFP966 and HDAC3 silencing increased WT-SIGIRR and IL-18Rα protein abundance without inducing their mRNA levels. HDAC3 inhibition delayed proteasome-dependent WT-SIGIRR turnover and stabilized IL-18Rα, thereby enhancing IL-37b-mediated anti-inflammatory signaling in CF airway epithelial cells.

## INTRODUCTION

Airway inflammation is a central feature of cystic fibrosis (CF) lung disease and persists despite major advances in CFTR modulator therapy.^1,2^ Exaggerated activation of innate immune pathways contributes to epithelial injury, mucus obstruction, infection, and progressive loss of lung function.^3,4^ However, the endogenous regulatory mechanisms that restrain mucosal inflammation in the CF airway remain incompletely understood. Defining pathways that restore epithelial immune homeostasis may therefore provide therapeutic opportunities that complement correction of the CFTR defect.

Single immunoglobulin IL-1 receptor-related molecule (SIGIRR), also known as TIR8 or IL-1R8, is a negative regulator of IL-1 receptor and Toll-like receptor (TLR) signaling.^5–7^ In addition to this canonical inhibitory function, SIGIRR is an essential component of the anti-inflammatory pathway mediated by interleukin-37 (IL-37), a member of the IL-1 cytokine family. IL-37b suppresses innate immune activation through a receptor complex containing SIGIRR and IL-18 receptor α (IL-18Rα).^8,9^ Disruption of this axis has been linked to excessive inflammatory responses in several disease contexts.^10,11^

SIGIRR function is also regulated by alternative splicing. An exon 8-skipped SIGIRR isoform (Δ8-SIGIRR) acts as a dominant-negative variant that interferes with plasma membrane localization and signaling competence of wild-type SIGIRR (WT-SIGIRR).^12,13^ In CF airway epithelial cells, increased Δ8-SIGIRR expression alters the balance between WT-SIGIRR and Δ8-SIGIRR, reduces cell-surface WT-SIGIRR, and limits IL-37-dependent anti-inflammatory activity.^14^ The molecular mechanisms that govern this imbalance, and the extent to which they can be therapeutically modulated, remain poorly defined.

Histone deacetylases (HDACs) are increasingly recognized as regulators of inflammatory gene expression, epithelial responses, and immune cell function.^15–17^ HDACs also modulate non-histone proteins, including factors involved in protein stability and trafficking.^18^ Whether HDAC activity controls the SIGIRR isoform balance or the IL-37 receptor module in CF airway epithelium has not been established.

Here, we identify HDAC3 as a regulator of WT-SIGIRR and IL-18Rα proteostasis in CF airway epithelial cells. We began with a transcript-based natural-product screen to identify compounds that preferentially enhance WT-SIGIRR expression over the dominant-negative Δ8-SIGIRR isoform. This screen identified short-chain fatty acids (SCFAs), endogenous microbial metabolites with HDAC-modulatory activity, as preferential enhancers of WT-SIGIRR. Subsequent mechanistic analyses using pan-HDAC inhibition, isoform-targeting pharmacological probes, HDAC3 silencing, protein stability assays, ubiquitination analysis, and primary CF airway epithelial cells revealed that HDAC3 regulates the IL-37 receptor module predominantly through receptor protein stability. Our findings, therefore, extend the regulation of IL-37b responsiveness from SIGIRR isoform expression to HDAC3-dependent receptor proteostasis and suggest that modulation of receptor stability may enhance endogenous anti-inflammatory signaling in CF airway epithelium.

## RESULTS

### SCFAs preferentially upregulate WT-SIGIRR over **Δ**8-SIGIRR

Membrane-associated SIGIRR is reduced in CF airway epithelial cells, at least in part because of increased expression of the dominant-negative Δ8-SIGIRR isoform. To distinguish WT-SIGIRR from Δ8-SIGIRR, we first established isoform-specific quantitative PCR assays using primer sets designed to selectively detect each transcript variant (Figure 1A). Using this assay system, we screened a library of 284 natural compounds in IB3-1 cells to identify compounds that preferentially increased WT-SIGIRR expression relative to Δ8-SIGIRR.

**Figure 1.**
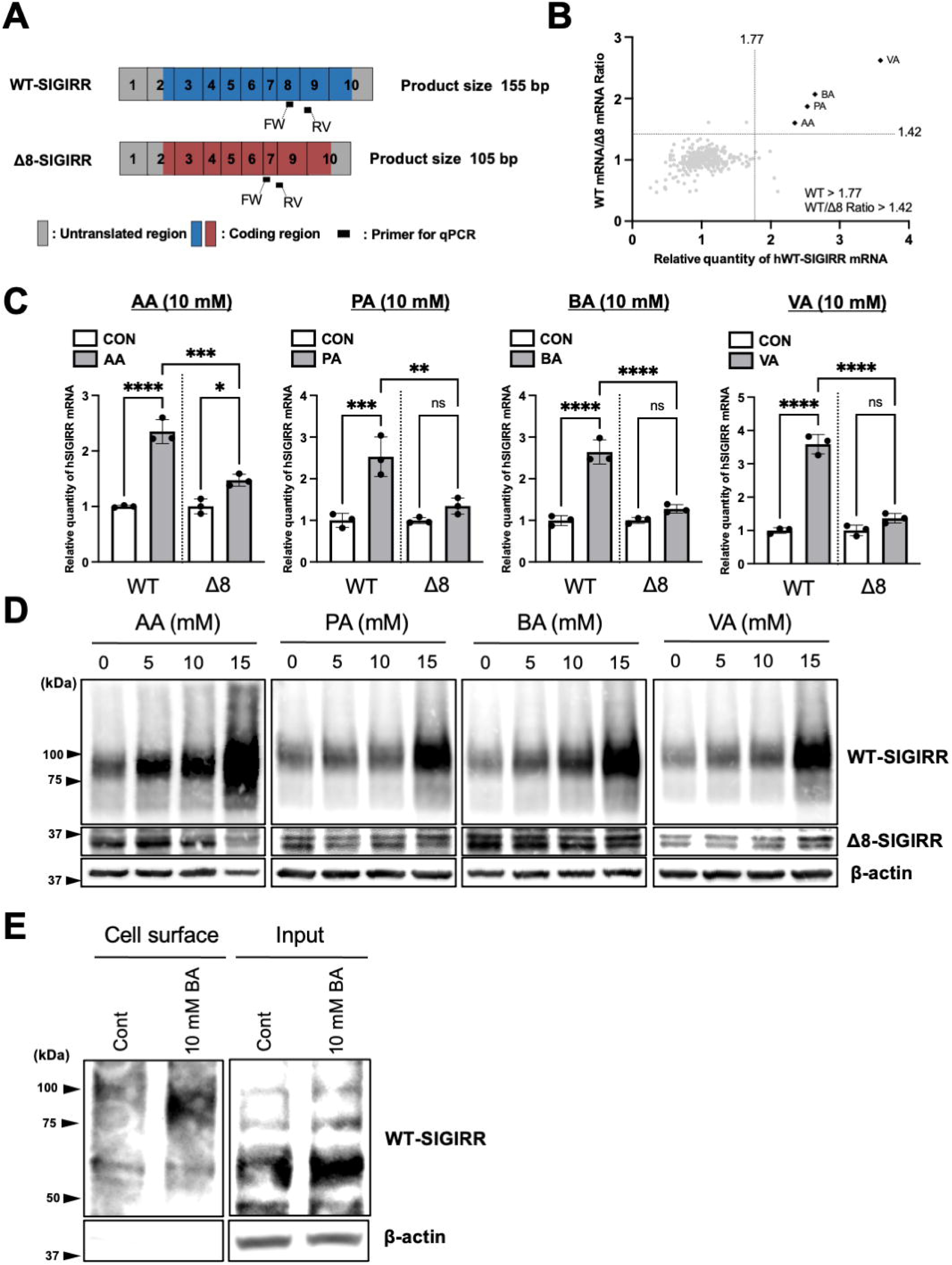
SCFAs preferentially upregulate WT-SIGIRR over Δ8-SIGIRR. (A) Schematic representation of the qPCR primer positions used to detect WT-SIGIRR and Δ8-SIGIRR. The reverse primer for Δ8-SIGIRR spans the exon 7–exon 9 junction. (B) A natural-compound library was screened in IB3-1 cells to identify compounds that preferentially increased WT-SIGIRR expression relative to Δ8-SIGIRR. The four hit compounds were short-chain fatty acids: acetic acid (AA), propionic acid (PA), butyric acid (BA), and valeric acid (VA). (C and D) IB3-1 cells were treated with the indicated concentrations of SCFAs for 24 h. WT-SIGIRR and Δ8-SIGIRR expression was analyzed by quantitative RT-PCR (C) and immunoblotting (D). Quantitative RT-PCR data were normalized to 18S rRNA. (E) Cell-surface WT-SIGIRR in IB3-1 cells treated with 10 mM BA for 24 h was assessed by biotinylation followed by immunoblotting. Data in (C) are presented as mean ± SD (n = 3). Statistical significance was assessed by one-way ANOVA followed by Tukey’s multiple-comparison test. *P* values are indicated in the graphs; ns, not significant; \**P* < 0.05; \*\**P* < 0.01; \*\*\**P* < 0.001; \*\*\*\**P* < 0.0001.

The compounds were ranked in descending order according to their induction levels of WT-SIGIRR mRNA expression (Figure S1). Nine compounds exceeded the screen-derived WT-SIGIRR induction cutoff value of 1.77, which was calculated from the 284-compound primary screening dataset as the mean relative WT-SIGIRR mRNA level plus two sample standard deviations. The four top-ranked compounds were all short-chain fatty acids (SCFAs): valeric acid (VA), butyric acid (BA), propionic acid (PA), and acetic acid (AA). We next evaluated selectivity for WT-SIGIRR by calculating the WT-SIGIRR/Δ8-SIGIRR mRNA ratio. The cutoff value for this ratio was similarly determined as the mean ratio plus two sample standard deviations, yielding a value of 1.42. Six compounds exceeded this ratio cutoff, including the same four SCFAs. Notably, these four SCFAs were the only compounds that satisfied both criteria: WT-SIGIRR induction above 1.77 and a WT/Δ8 induction ratio above 1.42 (Figure 1B). We then validated the effects of these SCFAs by qRT-PCR. All four SCFAs significantly increased WT-SIGIRR mRNA expression. Among them, AA also significantly increased Δ8-SIGIRR mRNA expression; however, the magnitude of Δ8-SIGIRR induction was substantially lower than that of WT-SIGIRR. In contrast, PA, BA, and VA did not significantly induce Δ8-SIGIRR expression (Figure 1C). These results indicate that SCFAs preferentially enhance WT-SIGIRR mRNA expression over Δ8-SIGIRR.

Consistent with the mRNA results, immunoblot analysis showed that SCFAs increased WT-SIGIRR protein abundance, particularly at higher concentrations. By contrast, Δ8-SIGIRR protein levels were not comparably increased, although a modest increase was observed after VA treatment (Figure 1D). Because functional SIGIRR is localized at the plasma membrane, we next examined cell-surface WT-SIGIRR expression. Cell-surface biotinylation analysis showed that BA increased WT-SIGIRR abundance at the cell surface, together with an increase in total WT-SIGIRR in the input fraction (Figure 1E).

Because SCFAs can alter extracellular pH, we further examined whether BA’s effect was attributable to nonspecific acidification. Treatment with sodium butyrate, the sodium salt of BA, also increased WT-SIGIRR protein abundance in a concentration-dependent manner (Figure S2). These results suggest that BA-dependent WT-SIGIRR upregulation is not simply attributable to nonspecific acidification. Together, these data indicate that SCFAs preferentially shift SIGIRR expression toward the functional WT isoform in CF airway epithelial cells.

### Pan-HDAC inhibition restores WT-SIGIRR expression and enhances IL-37b responses

Because SCFAs can signal through G protein-coupled receptors and can also inhibit HDAC activity,^19,20^ we next examined which pathway was associated with SCFA-dependent WT-SIGIRR induction. Pretreatment with pertussis toxin, an inhibitor of Gi/o-coupled receptor signaling, or YM-254890, an inhibitor of Gq/11-mediated signaling, did not attenuate SCFA-induced WT-SIGIRR mRNA upregulation (Figure S3). These results suggest that canonical G protein-dependent SCFA receptor signaling is not required for WT-SIGIRR induction under these conditions.

We therefore examined the effect of HDAC inhibition. The pan-HDAC inhibitor panobinostat, a clinically advanced broad HDAC inhibitor,^21,22^ preferentially increased WT-SIGIRR, but not Δ8-SIGIRR, at both the mRNA and protein levels (Figures 2A and 2B). Absolute transcript quantification by ddPCR confirmed that panobinostat increased WT-SIGIRR and reduced Δ8-SIGIRR transcript abundance, whereas BA increased WT-SIGIRR transcripts without increasing Δ8-SIGIRR (Figures 2C and 2D). We then tested whether panobinostat affected IL-37b responsiveness. IL-37b suppressed poly(I:C)-induced IL-8 production after panobinostat pretreatment, whereas IL-37b had little effect in the absence of panobinostat (Figure 2E). Thus, broad HDAC inhibition restores a WT-SIGIRR-favorable isoform balance and enables IL-37b-mediated anti-inflammatory signaling.

**Figure 2.**
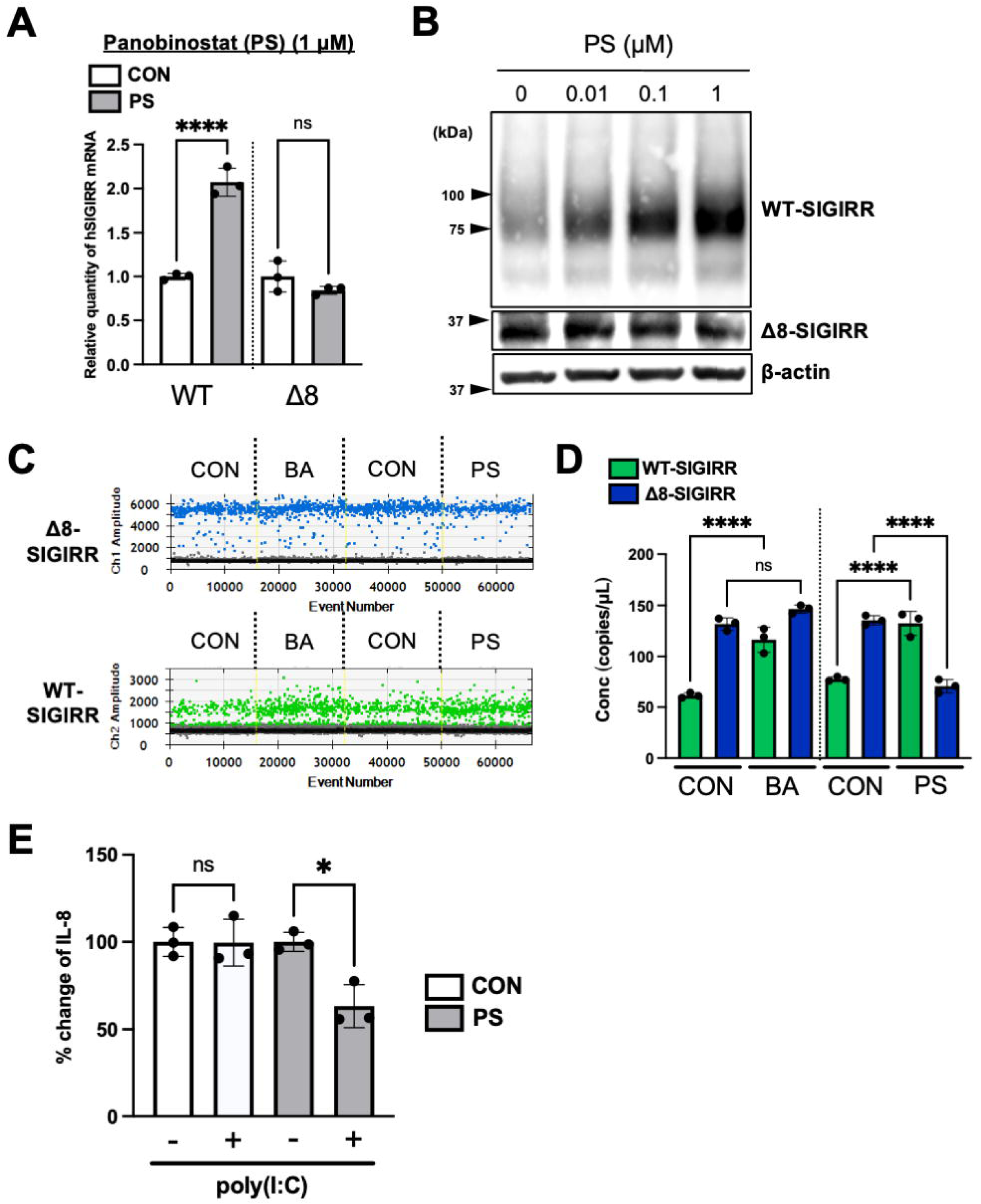
Pan-HDAC inhibition restores IL-37b-dependent anti-inflammatory activity via WT-SIGIRR upregulation. (A and B) IB3-1 cells were treated with the indicated concentrations of panobinostat (PS) for 24 h. WT-SIGIRR and Δ8-SIGIRR expression were analyzed by quantitative RT-PCR (A) and immunoblotting (B). Quantitative RT-PCR data were normalized to 18S rRNA. (C and D) IB3-1 cells were treated with 10 mM BA or 1 μM PS for 24 h. WT-SIGIRR and Δ8-SIGIRR transcripts were quantified by ddPCR. Representative droplet plots are shown in (C), and target concentrations are shown in (D). Quantification is presented as copies/μL and was calculated using QuantaSoft based on Poisson statistics. (E) IB3-1 cells were treated with DMSO or PS for 24 h under serum-free conditions. Cells were then treated with IL-37b for 2 h and stimulated with poly(I:C). After 24 h, IL-8 levels in culture supernatants were quantified by ELISA. Data in (A), (D), and (E) are presented as mean ± SD (n = 3). Statistical significance was assessed by one-way ANOVA followed by Tukey’s multiple-comparison test. *P* values are indicated in the graphs; ns, not significant; \**P* < 0.05; \*\*\*\**P* < 0.0001.

### HDAC3 inhibitor increases WT-SIGIRR mRNA only at high concentrations

HDACs comprise 18 isoforms grouped into class I (HDAC1, 2, 3, and 8), class II (HDAC4, 5, 6, 7, 9, and 10), class III (SIRT1–7), and class IV (HDAC11) enzymes, each with distinct roles in transcriptional and inflammatory regulation.^16,23^ To identify the HDAC isoform or isoforms involved in WT-SIGIRR regulation, IB3-1 cells were treated with isoform-selective inhibitors targeting HDAC1, 2, 3, 4, 6, 8, and 11 across a broad concentration range. The inhibitor panel included HDAC1-associated compound (-)-parthenolide, HDAC2 inhibitor santacruzamate A, HDAC3 inhibitor RGFP966, HDAC4 allosteric modulator tasquinimod, HDAC6 inhibitor CAY10603, HDAC8 inhibitor PCI34051, and HDAC11 inhibitor SIS17.^24–30^ Among the inhibitors tested, RGFP966 increased WT-SIGIRR mRNA, whereas Δ8-SIGIRR mRNA remained unchanged. This transcriptional effect was observed only at the highest concentration tested (10 μM), and no induction of WT-SIGIRR mRNA was detected at concentrations of 1 μM or lower (Figure 3C). Thus, pharmacological HDAC3 inhibition increased WT-SIGIRR mRNA only at high RGFP966 concentrations.

**Figure 3.**
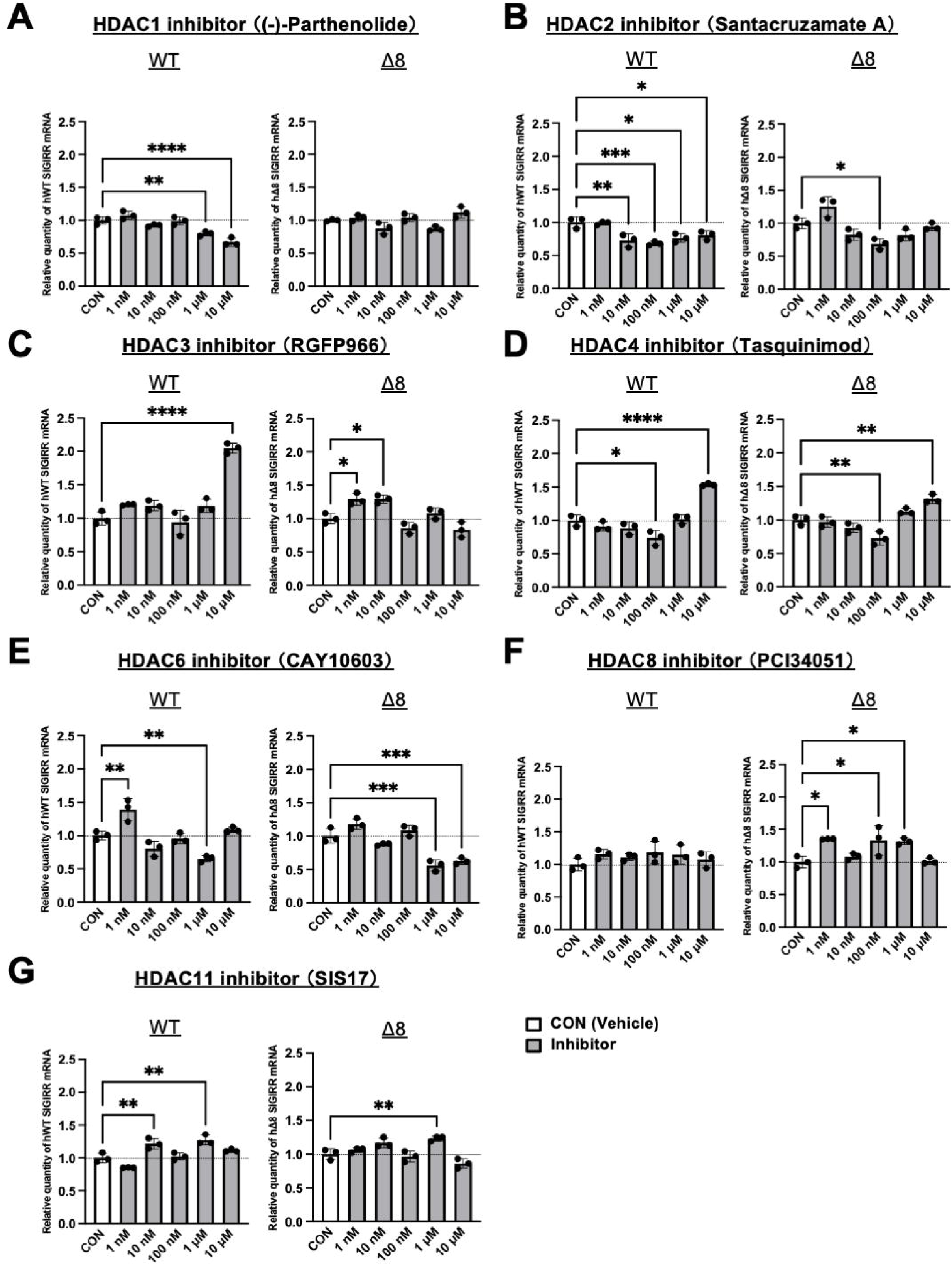
HDAC isoform-selective inhibitor screening identifies RGFP966 as an inducer of WT-SIGIRR mRNA expression. (A–G) IB3-1 cells were treated for 24 h with the indicated concentrations of HDAC isoform-selective inhibitors: the HDAC1 inhibitor-associated compound (-)-parthenolide (A), the HDAC2 inhibitor santacruzamate A (B), the HDAC3 inhibitor RGFP966 (C), the HDAC4 allosteric modulator tasquinimod (D), the HDAC6 inhibitor CAY10603 (E), the HDAC8 inhibitor PCI34051 (F), or the HDAC11 inhibitor SIS17 (G). WT-SIGIRR and Δ8-SIGIRR mRNA expression was analyzed by quantitative RT-PCR and normalized to 18S rRNA. Data are presented as mean ± SD (n = 3). Statistical significance was assessed by one-way ANOVA followed by Tukey’s multiple-comparison test. *P* values are indicated in the graphs; ns, not significant; \**P* < 0.05; \*\**P* < 0.01; \*\*\**P* < 0.001; \*\*\*\**P* < 0.0001.

### Low-concentration RGFP966 increases WT-SIGIRR and IL-18R**α** protein abundance without corresponding mRNA induction

We next examined RGFP966-dependent protein regulation across a broader concentration range (1 nM to 10 μM) in IB3-1 cells. WT-SIGIRR protein increased in a concentration-dependent manner beginning at 10 nM RGFP966, whereas Δ8-SIGIRR protein remained unchanged. IL-18Rα protein level increased in parallel (Figure 4A). At 1 μM RGFP966, WT-SIGIRR and IL-18Rα proteins increased without corresponding mRNA induction; at 10 μM, both protein and mRNA levels increased (Figure 4B). HDAC3 knockdown similarly increased WT-SIGIRR and IL-18Rα proteins (Figure 4C), whereas WT-SIGIRR mRNA was unchanged and IL-18Rα mRNA was reduced (Figure 4D). Comparable results were obtained in primary D-HBE-CF cells treated with RGFP966 or transfected with HDAC3 siRNA (Figures 4E–4H). These findings identify HDAC3 as a regulator of the IL-37 receptor module and indicate that post-transcriptional mechanisms predominate at low RGFP966 concentrations.

**Figure 4.**
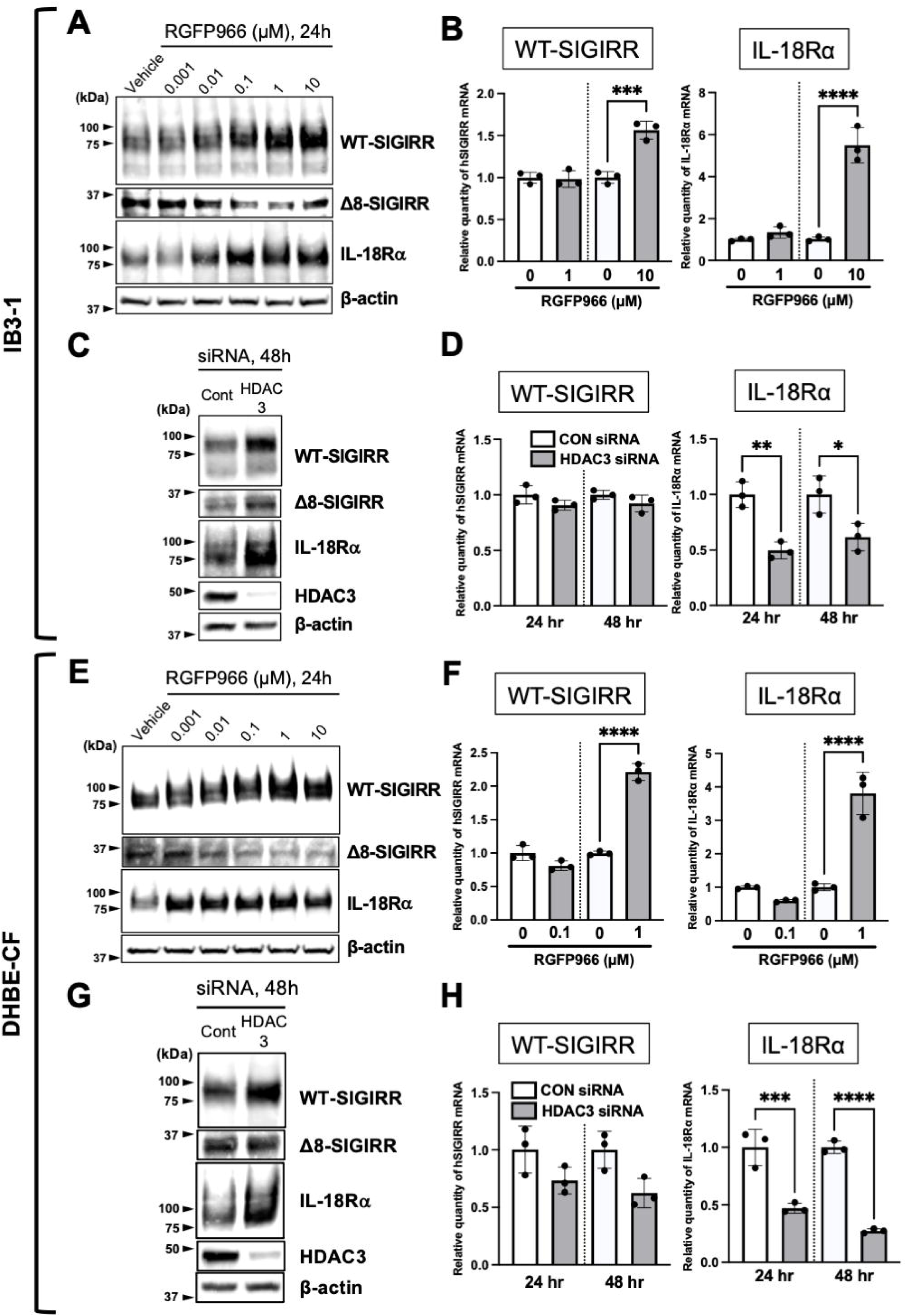
Low-concentration RGFP966 or HDAC3 knockdown increases WT-SIGIRR and IL-18Rα protein abundance without corresponding mRNA induction. (A and E) IB3-1 and D-HBE-CF cells were treated with the indicated concentrations of RGFP966 for 24 h, and target proteins were analyzed by immunoblotting. (B and F) IB3-1 and D-HBE-CF cells were treated with the indicated concentrations of RGFP966 for 24 h, and WT-SIGIRR and IL-18Rα mRNA expression was analyzed by quantitative RT-PCR. Quantitative RT-PCR data were normalized to 18S rRNA. (C and G) IB3-1 and D-HBE-CF cells were transfected with 20 nM non-targeting control siRNA or HDAC3 siRNA for 48 h, and target proteins were analyzed by immunoblotting. (D and H) IB3-1 and D-HBE-CF cells were transfected with 20 nM non-targeting control siRNA or HDAC3 siRNA for 24 or 48 h, and WT-SIGIRR and IL-18Rα mRNA expression was analyzed by quantitative RT-PCR. Quantitative RT-PCR data were normalized to 18S rRNA. Data in (B), (D), (F), and (H) are presented as mean ± SD (n = 3). Statistical significance was assessed by one-way ANOVA followed by Tukey’s multiple-comparison test. *P* values are indicated in the graphs; ns, not significant; \**P* < 0.05; \*\**P* < 0.01; \*\*\**P* < 0.001; \*\*\*\**P* < 0.0001.

### HDAC3 inhibition stabilizes WT-SIGIRR and IL-18R**α** through a proteasome-dependent mechanism

To determine whether increased receptor abundance reflected altered protein stability, we performed cycloheximide chase experiments. In IB3-1 cells, 1 μM RGFP966 delayed the degradation of both WT-SIGIRR and IL-18Rα (Figure 5A). A similar stabilization effect was observed at 10 μM RGFP966 (Figure S4), indicating that protein stabilization contributes to receptor upregulation even at concentrations that also increase WT-SIGIRR and IL-18Rα mRNA. HDAC3 knockdown also delayed degradation of WT-SIGIRR and IL-18Rα (Figure 5B). To determine the degradation pathway, cells were treated with the proteasome inhibitor MG132 or the lysosomal inhibitor chloroquine. MG132 increased WT-SIGIRR and IL-18Rα protein abundance (Figure 5C), whereas chloroquine induced LC3 accumulation without increasing WT-SIGIRR or IL-18Rα (Figure 5D). These data support proteasome-dependent turnover of the receptor module.

**Figure 5.**
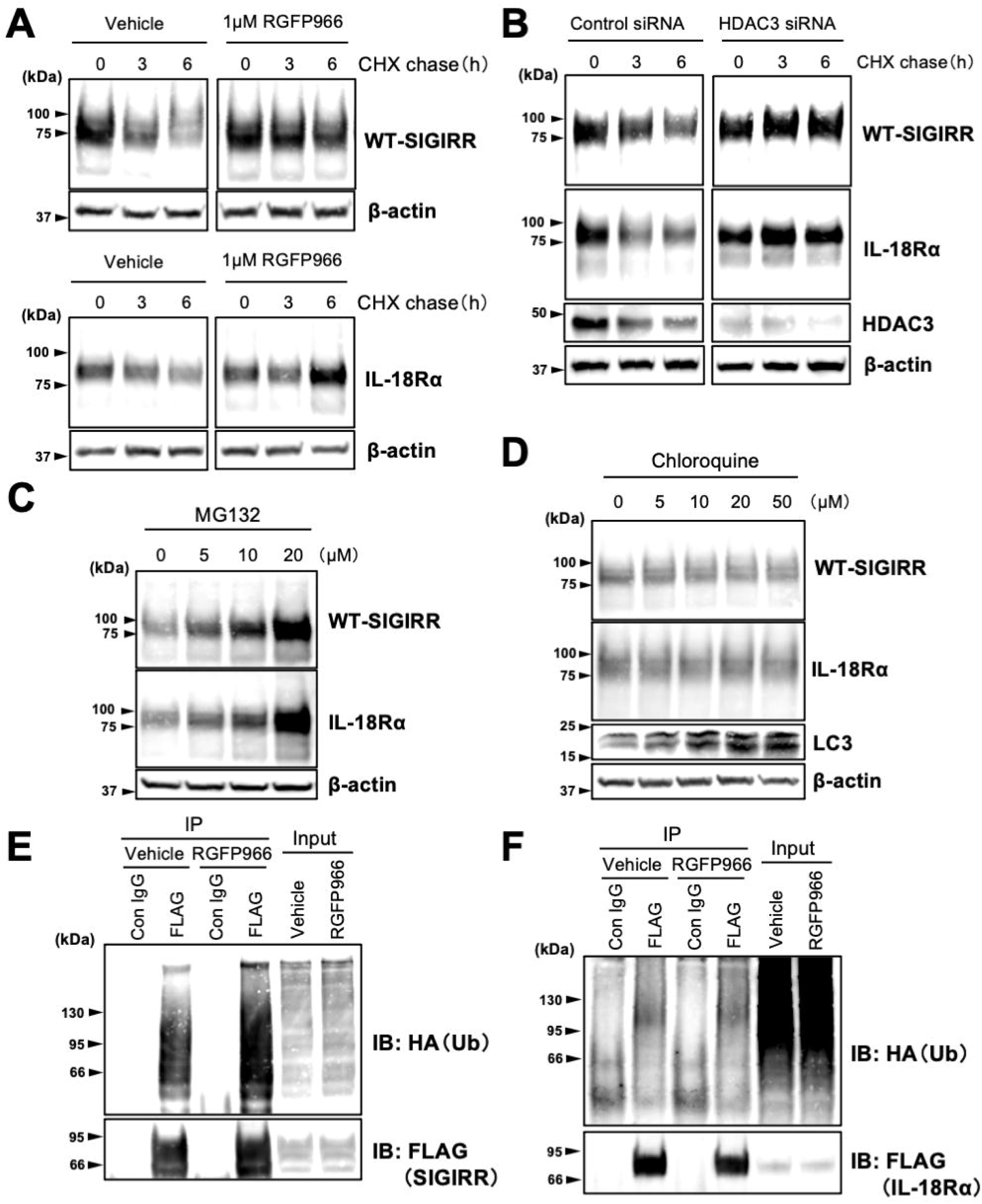
HDAC3 inhibition stabilizes WT-SIGIRR and IL-18Rα through a proteasome-dependent mechanism. (A) IB3-1 cells were treated with cycloheximide (CHX) to inhibit de novo protein synthesis in the presence or absence of 1 μM RGFP966. Cells were harvested at the indicated time points, and target proteins were analyzed by immunoblotting. (B) IB3-1 cells were transfected with 20 nM non-targeting control siRNA or HDAC3 siRNA for 48 h and then treated with CHX. Cells were harvested at the indicated time points, and target proteins were analyzed by immunoblotting. (C and D) IB3-1 cells were treated with the indicated concentrations of MG132 or chloroquine for 6 h, and target proteins were analyzed by immunoblotting. (E and F) IB3-1 cells were co-transfected with WT-SIGIRR-FLAG or IL-18Rα-FLAG and HA-tagged ubiquitin. Twenty-four hours after transfection, the cells were treated with 1 μM RGFP966 for 24 h. Whole-cell lysates were subjected to immunoprecipitation with an anti-FLAG antibody, and ubiquitinated WT-SIGIRR or IL-18Rα was detected by immunoblotting with an anti-HA antibody. Immunoprecipitated WT-SIGIRR or IL-18Rα was confirmed by immunoblotting with an anti-FLAG antibody. IgG immunoprecipitation was included as a negative control. Representative blots are shown.

Ubiquitination assays revealed an accumulation of ubiquitin-positive WT-SIGIRR species after RGFP966 treatment (Figure 5E). Together with the cycloheximide chase and MG132 experiments, this finding is consistent with reduced clearance or delayed proteasomal processing of ubiquitinated WT-SIGIRR. In contrast, ubiquitin-positive IL-18Rα species did not increase detectably under the same conditions (Figure 5F), suggesting that IL-18R α stabilization may occur indirectly or through ubiquitin modifications that are not efficiently captured by this assay. Because lysine acetylation can modulate protein ubiquitination and stability, we next examined whether RGFP966 increased lysine acetylation of WT-SIGIRR. Immunoprecipitation of FLAG-tagged WT-SIGIRR followed by immunoblotting with an anti-acetyl-lysine antibody did not reveal a detectable increase in WT-SIGIRR acetylation after RGFP966 treatment (Figure S5). Thus, the stabilization of WT-SIGIRR by HDAC3 inhibition does not appear to be explained by a detectable increase in bulk lysine acetylation of WT-SIGIRR itself. Instead, HDAC3 inhibition may indirectly regulate SIGIRR proteostasis, potentially through effects on ubiquitin-processing enzymes, chaperones, trafficking factors, or other components of the proteasome-dependent degradation machinery.

### HDAC3 inhibition enhances IL-37b-mediated suppression of inflammatory responses

Finally, we examined whether stabilization of WT-SIGIRR and IL-18Rα enhanced IL-37b responsiveness. In IB3-1 cells, pretreatment with RGFP966 (1 μM) enhanced the suppressive effect of IL-37b on poly(I:C)-induced IL-8 production (Figure 6A). A similar enhancement was observed following HDAC3 knockdown, indicating that this effect was associated with HDAC3 inhibition (Figure 6B). In primary D-HBE-CF cells, low-concentration RGFP966 (0.1 μM) also augmented IL-37b-mediated suppression of IL-8 production induced by poly(I:C) or the TLR5 ligand *Pseudomonas aeruginosa* flagellin (Figures 6C and 6D). Collectively, these results indicate that HDAC3 inhibition enhances IL-37b-mediated anti-inflammatory signaling in CF airway epithelial cells, at least in part by stabilizing key components of the WT-SIGIRR/IL-18Rα receptor module (Figure 6E).

**Figure 6.**
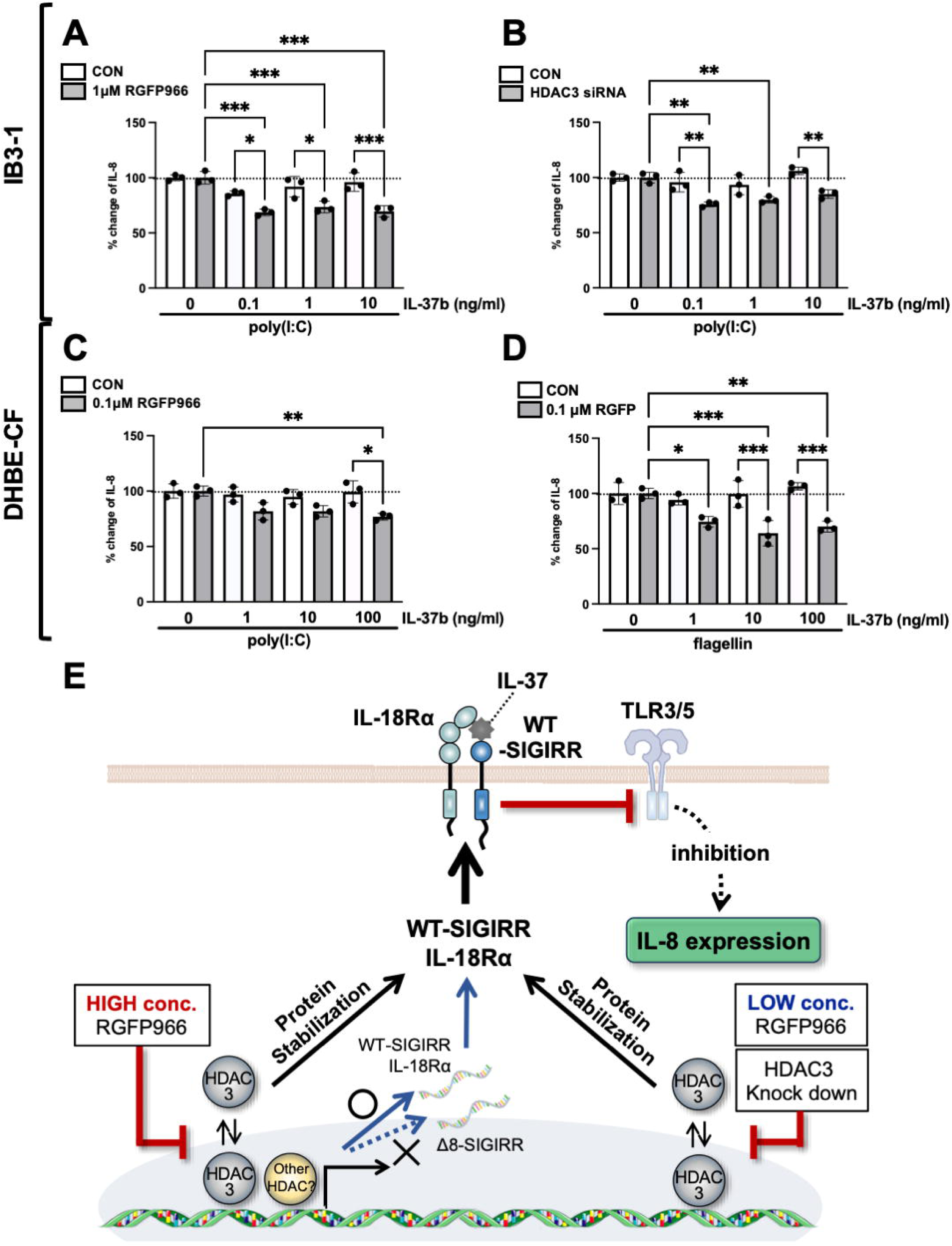
HDAC3 inhibition enhances IL-37b-mediated anti-inflammatory activity. (A) IB3-1 cells were treated with 1 μM RGFP966 for 24 h and then treated with the indicated concentrations of IL-37b. Cells were stimulated with 20 μg/mL poly(I:C) 2 h later. After 24 h, IL-8 levels in culture supernatants were quantified by ELISA. (B) IB3-1 cells were transfected with 20 nM non-targeting control siRNA or HDAC3 siRNA for 48 h and then treated with the indicated concentrations of IL-37b. Cells were stimulated with 20 μg/mL poly(I:C) 2 h later, and IL-8 levels were measured after 24 h. (C) D-HBE-CF cells were treated with 0.1 μM RGFP966 for 24 h and then treated with the indicated concentrations of IL-37b. Cells were stimulated with 500 ng/mL poly(I:C) 2 h later, and IL-8 levels were measured after 24 h. (D) D-HBE-CF cells were treated with 0.1 μM RGFP966 for 24 h and then treated with the indicated concentrations of IL-37b. Cells were stimulated with 100 ng/mL flagellin 2 h later, and IL-8 levels were measured after 24 h. Data in (A)–(D) are presented as mean ± SD (n = 3). Statistical significance was assessed by one-way ANOVA followed by Tukey’s multiple-comparison test. *P* values are indicated in the graphs; ns, not significant; \**P* < 0.05; \*\**P* < 0.01; \*\*\**P* < 0.001. (E) Proposed model for HDAC3-dependent regulation of the IL-37 receptor module in CF airway epithelial cells. HDAC3 inhibition increases the abundance of WT-SIGIRR and IL-18Rα proteins, thereby enhancing IL-37b-dependent anti-inflammatory signaling. This mechanism is primarily associated with stabilization of WT-SIGIRR and IL-18Rα proteins and may be accompanied by transcriptional upregulation of WT-SIGIRR at high RGFP966 concentrations.

## DISCUSSION

SIGIRR is a critical negative regulator of IL-1 receptor and TLR signaling and, together with IL-18Rα, forms an essential receptor module for IL-37b-mediated anti-inflammatory activity. In CF airway epithelial cells, increased expression of the dominant-negative Δ8-SIGIRR isoform limits WT-SIGIRR membrane localization and impairs IL-37-dependent suppression of inflammatory signaling. The present study identifies HDAC3 as a previously unrecognized regulator of this pathway and supports a model in which HDAC3 constrains the availability of the WT-SIGIRR/IL-18Rα receptor module through both transcriptional and post-transcriptional mechanisms. As summarized in Figure 6E, low-concentration RGFP966 and HDAC3 knockdown primarily enhance receptor protein stability, whereas high-concentration RGFP966 additionally promotes WT-SIGIRR mRNA induction. Together, these effects increase the functional IL-37b receptor module and strengthen epithelial suppression of poly(I:C)-driven inflammatory responses. An important conceptual point emerging from this study is that a transcript-based discovery strategy ultimately uncovered a protein-level mechanism controlling the IL-37 receptor module. The initial screen was designed to identify compounds that preferentially increased WT-SIGIRR mRNA expression relative to Δ8-SIGIRR. However, subsequent analyses showed that low-concentration RGFP966 and HDAC3 knockdown increased WT-SIGIRR and IL-18Rα primarily by stabilizing receptor proteins rather than by increasing their mRNA levels. Thus, the study progressed from SIGIRR isoform-focused transcriptional screening to the identification of HDAC3-dependent receptor proteostasis as a regulatory layer of IL-37b-mediated anti-inflammatory signaling.

A key finding of this study is that SCFAs shift SIGIRR regulation toward the functional WT isoform and preferentially enhance WT-SIGIRR expression over Δ8-SIGIRR. This is consistent with previous reports linking SCFAs to SIGIRR induction and epithelial immune regulation.^19,20,31^ Because SCFAs are microbial metabolites that can influence mucosal immune homeostasis, this observation suggests that metabolite-dependent regulation may help maintain or restore epithelial anti-inflammatory pathways. The ability of sodium butyrate to reproduce BA-dependent SIGIRR receptor upregulation argues against a simple acidification-related mechanism. In addition, inhibition of canonical Gi/o- or Gq/11-dependent signaling did not attenuate SCFA-induced WT-SIGIRR expression. Although these experiments do not exclude all receptor-dependent or noncanonical SCFA signaling mechanisms, they support the interpretation that the observed response is not primarily driven by classical G protein-dependent SCFA receptor signaling under the conditions tested. These observations provided the rationale for focusing on HDAC-dependent regulation.

Pan-HDAC inhibition corrected the SIGIRR isoform imbalance and restored IL-37b responsiveness, indicating that HDAC activity contributes to repression or destabilization of the IL-37 receptor axis in CF airway epithelial cells. However, pan-HDAC inhibitors affect multiple HDAC isoforms and numerous transcriptional and non-transcriptional pathways. Therefore, identifying HDAC3 as a principal regulator is important because it provides a more precise entry point into the mechanism that controls receptor availability. The combined pharmacological and genetic data support a model in which HDAC3 primarily regulates WT-SIGIRR and IL-18Rα at the post-transcriptional level, whereas broader transcriptional effects may emerge at higher inhibitor concentrations.

The concentration-dependent behavior of RGFP966 provides an important framework for interpreting HDAC3-dependent regulation. Low concentrations of RGFP966 increased WT-SIGIRR and IL-18Rα protein levels without a corresponding increase in mRNA, whereas high concentrations also increased WT-SIGIRR mRNA. Because high-concentration RGFP966 also delayed receptor degradation, receptor upregulation at this concentration is best interpreted as the combined consequence of transcriptional induction and enhanced protein stability. In contrast, the effects of low-concentration RGFP966 and HDAC3 knockdown point to a predominantly post-transcriptional mechanism. This distinction is important, as RGFP966 is widely used as an HDAC3-selective inhibitor but has been reported to exhibit slow binding and measurable activity toward HDAC1 and HDAC2 under certain conditions.^32^ Thus, the low-concentration and knockdown data provide the strongest evidence for HDAC3-dependent proteostatic regulation, whereas high-concentration transcriptional effects should be interpreted as potentially involving broader class I HDAC activity.

Mechanistically, our data implicate ubiquitin-proteasome-dependent turnover as a major determinant of WT-SIGIRR and IL-18Rα availability. The increase in WT-SIGIRR protein abundance after RGFP966 treatment or HDAC3 knockdown, together with delayed degradation in cycloheximide chase experiments and receptor accumulation after MG132 treatment, supports a model in which WT-SIGIRR is normally limited by proteasome-dependent turnover. Importantly, RGFP966 treatment led to the accumulation of ubiquitin-positive WT-SIGIRR species. This finding should not be interpreted simply as enhanced ubiquitination that promotes degradation. Rather, in the context of increased WT-SIGIRR stability, it is more consistent with slowed clearance or delayed proteasomal processing of ubiquitinated WT-SIGIRR. Thus, HDAC3 inhibition may affect one or more steps downstream of ubiquitin attachment, including recognition, extraction, trafficking, deubiquitination, or proteasomal processing of ubiquitinated receptor species. This interpretation is consistent with prior evidence that the deubiquitinase USP13 stabilizes IL-1R8/SIGIRR and suppresses lung inflammation, further supporting the concept that ubiquitin-dependent proteostasis is central to controlling SIGIRR function in airway inflammatory responses.^33^

IL-18Rα regulation appears to be more complex. Although IL-18Rα was stabilized together with WT-SIGIRR, ubiquitin-positive IL-18Rα species did not detectably accumulate under the present conditions. One possibility is that IL-18Rα is indirectly stabilized through improved assembly, trafficking, or membrane retention of the WT-SIGIRR/IL-18Rα receptor module. Receptor complex formation can influence membrane protein turnover, intracellular trafficking, and cell-surface retention, providing a plausible framework in which stabilization of WT-SIGIRR secondarily enhances IL-18Rα stability.^34^ Alternatively, IL-18Rα ubiquitination may occur at low abundance, at specific lysine residues, or through linkage types that were not efficiently detected by the current assay. Defining whether IL-18Rα is directly regulated by HDAC3-dependent proteostasis or secondarily stabilized through WT-SIGIRR-containing receptor complexes will be an important area for future study.

Protein acetylation provides a plausible mechanistic link between HDAC activity and ubiquitin-dependent proteostasis. Lysine acetylation can influence ubiquitination, protein turnover, and the fate of non-histone proteins, and HDAC-dependent regulation of protein stability has been described in multiple contexts.^35,36^ In particular, HDAC3 has been reported to regulate the stability or signaling activity of several non-histone proteins through pathways connected to proteasome-mediated degradation, including cyclin A, Notch pathway components, and p21.^37,38, 39^ In addition, recent studies further illustrate how acetylation-ubiquitination crosstalk can regulate protein stability in disease-relevant signaling pathways.^40^ The intracellular region of SIGIRR contains lysine residues that could potentially serve as regulatory sites for acetylation or ubiquitination.^41^ However, WT-SIGIRR lysine acetylation was not detectably increased after RGFP966 treatment using the present immunoprecipitation-based approach. This finding argues against a model in which HDAC3 inhibition stabilizes WT-SIGIRR through a readily detectable increase in bulk lysine acetylation of WT-SIGIRR itself. Nevertheless, transient, low-stoichiometry, or site-specific acetylation cannot be excluded. An alternative model is that HDAC3 regulates upstream components of the receptor proteostasis machinery, such as E3 ubiquitin ligases, deubiquitinases, chaperones, trafficking regulators, or proteasome-associated factors, thereby controlling the fate of ubiquitinated WT-SIGIRR without requiring detectable acetylation of WT-SIGIRR itself.

Functionally, HDAC3 inhibition enhanced IL-37b-mediated suppression of poly(I:C) and *P. aeruginosa* flagellin-triggered epithelial inflammatory responses. This is relevant to CF airway disease, in which viral products and *P. aeruginosa* flagellin can amplify epithelial production of neutrophil-recruiting mediators such as IL-8. The model emerging from this study is that HDAC3 inhibition increases the availability of WT-SIGIRR and IL-18Rα at the protein level, thereby enhancing IL-37b’s capacity to restrain TLR-driven inflammatory output. The use of both IB3-1 cells and primary CF airway epithelial cells supports the disease relevance of this pathway, although additional validation in multiple donor-derived primary cultures and differentiated air-liquid interface models will be required to determine its broader applicability.

In summary, this study identifies HDAC3-dependent receptor proteostasis as a mechanism that limits the IL-37b-WT-SIGIRR/IL-18Rα anti-inflammatory axis in CF airway epithelial cells. Although the study began with an mRNA-based screen for compounds that preferentially enhance WT-SIGIRR expression, the subsequent mechanistic analyses revealed that receptor protein stability is a major determinant of IL-37b responsiveness. SCFAs and broad HDAC inhibition shift SIGIRR regulation toward the functional WT isoform, while HDAC3 inhibition stabilizes key IL-37 receptor components and enhances epithelial anti-inflammatory responsiveness. The proposed model highlights a concentration-dependent distinction: high-concentration RGFP966 can induce WT-SIGIRR transcription in addition to stabilizing receptor proteins, whereas low-concentration RGFP966 and HDAC3 knockdown primarily stabilize receptor proteins. These findings reveal receptor proteostasis as a regulatory layer of mucosal innate immune control and suggest that modulation of receptor stability may provide a strategy to restore endogenous anti-inflammatory capacity in CF airway disease.

### Limitations of the study

Several limitations should be considered. The molecular intermediates connecting HDAC3 to WT-SIGIRR and IL-18Rα turnover remain unidentified. The present study does not define the ubiquitin linkage type, ubiquitination sites, or specific degradation machinery responsible for WT-SIGIRR turnover. The mechanism underlying IL-18Rα co-stabilization also remains unresolved. In addition, although sodium butyrate and GPCR inhibitor experiments help exclude simple acidification and canonical G protein-dependent SCFA receptor signaling as dominant explanations, they do not fully define the SCFA-responsive pathway. Finally, the functional analysis focused primarily on IL-8 secretion as an inflammatory readout. Future studies incorporating receptor-loss and rescue experiments, HDAC3 rescue with wild-type and catalytically inactive mutants, ubiquitination-site mapping, identification of the relevant E3 ligase or deubiquitinase, broader cytokine profiling, and differentiated airway epithelial models will be needed to establish the full scope and mechanism of HDAC3-dependent regulation.

## Supporting information

Source Data uncropped blots

## RESOURCE AVAILABILITY

### Lead Contact

Further information and requests for resources and reagents should be directed to, and will be fulfilled by, the lead contact: Tsuyoshi Shuto.

### Materials availability

Plasmids and other unique reagents generated in this study are available from the lead contact upon reasonable request, subject to institutional and material-transfer restrictions. The natural-product library used in this study is available, subject to institutional policies and compound availability.

### Data and code availability

- This paper does not report original code.
- All data reported in this paper are included in the manuscript and supplemental information.
- Any additional information required to reanalyze the data reported in this paper is available from the lead contact upon request.

## ACKNOWLEDGMENTS

This work was supported by Japan Society for the Promotion of Science (JSPS) KAKENHI grants JP22K06642 to K.U.-S. and JP23K06150 to T.S.; the Program for Building Regional Innovation Ecosystems at Kumamoto University; and the Program for Fostering Innovators to Lead a Better Co-being Society (JPMJSP2127; MEXT, Japan). The authors thank the members of the Department of Molecular Medicine, Graduate School of Pharmaceutical Sciences, Kumamoto University, for technical assistance and helpful discussions.

## AUTHOR CONTRIBUTIONS

Conceptualization, K.U.-S. and T.S.; Methodology, K.U.-S., A.F., K.N., Y.H., S.T., and T.S.; Investigation, K.U.-S., A.F., K.N., T.K., K.K., and T.S.; Formal Analysis, K.U.-S., A.F., K.N., T.K., K.K., and T.S.; Resources, Y.H. and S.T.; Writing – Original Draft, K.U.-S., M.A.S., and T.S.; Writing – Review & Editing, K.U.-S., A.F., K.N., Y.H., S.T., T.K., K.K., M.A.S., and T.S.; Supervision, T.S.; Funding Acquisition, K.U.-S. and T.S.

## SUPPLEMENTAL INFORMATION

Document S1. Figures S1–S5, uncropped blots

## Supplemental figure legends

**Figure S1. Screening of natural compounds that induce WT-SIGIRR and Δ8-SIGIRR mRNA expression in IB3-1 cells**

IB3-1 cells were treated with 284 natural compounds for 24 h, and the relative mRNA expression levels of WT-SIGIRR and Δ8-SIGIRR were evaluated by quantitative RT-PCR. Compounds are ranked in descending order according to the induction level of WT-SIGIRR mRNA. Red bars indicate WT-SIGIRR mRNA expression, and blue bars indicate Δ8-SIGIRR mRNA expression. Values are shown as relative quantities normalized to the control condition. The inset shows an enlarged view of the top four WT-SIGIRR-inducing compounds. The top-ranked compounds were all short-chain fatty acids: 1, VA, valeric acid; 2, BA, butyric acid; 3, PA, propionic acid; and 4, AA, acetic acid.

**Figure S2. Sodium butyrate increases WT-SIGIRR protein abundance**

IB3-1 cells were treated with the indicated concentrations of sodium butyrate, the sodium salt of butyric acid, for 24 h. WT-SIGIRR protein levels were analyzed by immunoblotting using an anti-SIGIRR antibody. β-actin was used as a loading control. Sodium butyrate increased WT-SIGIRR protein abundance, suggesting that BA-induced receptor upregulation is not simply attributable to nonspecific acidification. Representative blots are shown.

**Figure S3. G protein-coupled receptor signaling inhibitors do not attenuate SCFA-induced WT-SIGIRR mRNA upregulation**

IB3-1 cells were pretreated for 1 h with 250 ng/mL pertussis toxin (PTX), an inhibitor of Gi/o-coupled receptor signaling, or 1 μM YM-254890, an inhibitor of Gq/11-mediated signaling. Cells were then treated with the 10 mM indicated SCFAs—acetic acid (AA), propionic acid (PA), butyric acid (BA), or valeric acid (VA)—for 24 h in the continued presence or absence of each inhibitor. WT-SIGIRR mRNA expression was analyzed by quantitative RT-PCR and normalized to 18S rRNA. Data are presented as mean ± SD (n = 3). Statistical significance was assessed by one-way ANOVA followed by Tukey’s multiple-comparison test. *P* values are indicated in the graphs; ns, not significant; \**P* < 0.05; \*\**P* < 0.01; \*\*\**P* < 0.001; \*\*\*\**P* < 0.0001.

**Figure S4. High-concentration RGFP966 stabilizes WT-SIGIRR and IL-18Rα during cycloheximide chase**

IB3-1 cells were treated with cycloheximide (CHX) to inhibit de novo protein synthesis in the presence or absence of 10 μM RGFP966. Cells were harvested at the indicated time points, and WT-SIGIRR and IL-18Rα protein levels were analyzed by immunoblotting. β-actin was used as a loading control. RGFP966 delayed the decline in WT-SIGIRR and IL-18Rα protein abundance after CHX treatment, supporting a protein-stabilizing component of RGFP966-induced receptor upregulation. Representative blots are shown.

Figure S5. RGFP966 does not detectably increase lysine acetylation of WT-SIGIRR IB3-1 cells expressing FLAG-tagged WT-SIGIRR were treated with 1 μM RGFP966 for 24 h. Whole-cell lysates were subjected to immunoprecipitation with an anti-FLAG antibody, and lysine acetylation of immunoprecipitated WT-SIGIRR was assessed by immunoblotting with an anti-acetyl-lysine antibody. Immunoprecipitated WT-SIGIRR was confirmed by immunoblotting with an anti-FLAG antibody. Control IgG immunoprecipitation was included as a negative control. RGFP966 did not produce a detectable increase in lysine acetylation of WT-SIGIRR under these conditions. Representative blots are shown.

## METHODS

Key resources table

2 Key resources table
| REAGENT or RESOURCE | SOURCE | IDENTIFIER |
| --- | --- | --- |
| Experimental models: cell lines |  |  |
| IB3-1 cells | ATCC | Cat#CRL-2777 |
| D-HBE-CF cells | Lonza | 38-year-old male Caucasian donor, lot No. 222158, with no reported history of smoking or alcohol use |
| Chemicals, peptides, and recombinant proteins |  |  |
| Acetic acid | Tokyo Chemical Industry (TCI, Tokyo, Japan) | Cat#A2035 |
| Propionic acid | Tokyo Chemical Industry (TCI, Tokyo, Japan) | Cat#P0500 |
| Butyric acid | Tokyo Chemical Industry (TCI, Tokyo, Japan) | Cat#B0754 |
| Valeric acid | Tokyo Chemical Industry (TCI, Tokyo, Japan) | Cat#V0003 |
| Panobinostat | Selleck Chemicals (Houston, TX, USA) | Cat#S1030 |
| SIS17 | Selleck Chemicals (Houston, TX, USA) | Cat#S6687 |
| Chloroquine | Selleck Chemicals (Houston, TX, USA) | Cat#S6999 |
| Tasquinimod | Cayman Chemical (Ann Arbor, MI, USA) | Cat#17692 |
| CAY10603 | Cayman Chemical (Ann Arbor, MI, USA) | Cat#13146 |
| (-)-Parthenolide | MedChemExpress (USA) | Cat#HY-N0141 |
| Santacruzamate A | MedChemExpress (USA) | Cat#HY-N0931 |
| RGFP966 | MedChemExpress (USA) | Cat#HY-13909 |
| PCI34051 | MedChemExpress (USA) | Cat#HY-15224 |
| Low-molecular-weight poly(I:C) | InvivoGen (San Diego, CA, USA) | Cat#tlrl-picw |
| Ultrapure *Pseudomonas aeruginosa* flagellin | InvivoGen (San Diego, CA, USA) | Cat#tlrl-pafla |
| Recombinant Val46 IL-37b | R&D Systems (Minneapolis, MN, USA) | Cat#7585-IL-025 |
| Cycloheximide | Wako (Osaka, Japan) | Cat#037-20991 |
| MG132 | Sigma-Aldrich (St. Louis, MO) | Cat#M7449 |
| Pertussis Toxin Solution | Wako (Osaka, Japan) | Cat#168-22471 |
| YM-254890 | Wako (Osaka, Japan) | Cat#257-00631 |
| Antibodies |  |  |
| Anti-SIGIRR antibody | R&D Systems (Minneapolis, MN, USA) | Cat#AF990<br>RRID:AB_218902<br>2 |
| Anti-IL-18Rα antibody | R&D Systems (Minneapolis, MN, USA) | Cat#AF840<br>RRID:AB_355650 |
| Anti-HDAC3 antibody | Proteintech (USA) | Cat#10255-1-AP<br>RRID:AB_227973<br>3 |
| Anti-LC3 antibody | Proteintech (USA) | Cat#14600-1-AP<br>RRID:AB_213773<br>7 |
| Anti-β-actin antibody | Bio-Rad Laboratories (USA) | Cat#MCA5775GA<br>RRID:AB_257158<br>0 |
| Anti-FLAG antibody | Wako (Osaka, Japan) | Cat#018-22381 |
| Anti-HA antibody | Proteintech (USA) | Cat#51064-2-AP<br>RRID:AB_110423<br>21 |
| Anti-pan acetylation antibody | Proteintech (USA) | Cat#66289-1-Ig<br>RRID:AB_288167<br>2 |
| Normal mouse IgG | Santa Cruz Biotechnology (USA) | Cat#sc-2025<br>RRID:AB_737182 |
| Recombinant DNA |  |  |
| pDNA3.1-HA-Ubiquitin | Bioworld Technology (MN, USA) | Cat#PPL00017-2e |
| pCMV-WT-SIGIRR-FLAG | This paper | Full-length human WT-SIGIRR cloned into pCMV-FLAG-C |
| Critical commercial assays |  |  |
| NucleoSpin RNA Plus | TaKaRa (Tokyo, Japan) | Cat#U0984B |
| PrimeScript RT Master Mix | TaKaRa (Tokyo, Japan) | Cat#RR036A |
| ddPCR Supermix for probes, no dUTP | Bio-Rad Laboratories (USA) | Cat#186-3023 |
| FuGENE HD | Promega (Madison, WA, USA) | Cat#E2311 |
| Lipofectamine RNAiMAX | Thermo Fisher Scientific (USA) | Cat#13778-030 |
| siGENOME non-targeting siRNA control pool | Dharmacon | Cat#D-001206-14-05 |
| siGENOME human HDAC3 siRNA SMARTpool | Dharmacon | Cat#M-003496-02-0005 |
| EZ-Link Sulfo-NHS-SS-Biotin | Thermo Fisher Scientific (USA) | Cat#21331 |
| Streptavidin agarose resin | Thermo Fisher Scientific (USA) | Cat#20347 |
| Protein A/G PLUS-Agarose | Santa Cruz Biotechnology (USA) | Cat#sc-2003 |
| DuoSet human IL-8 ELISA kit | R&D Systems (Minneapolis, MN, USA) | Cat#DY208 |
| Software and algorithms |  |  |
| GraphPad Prism 11 | GraphPad Software (San Diego, CA, USA) | Version 11 |
| QuantaSoft | Bio-Rad Laboratories (USA) | Version 1.7.4.0917 |
| Other |  |  |
| QX200 AutoDG Droplet Digital PCR system | Bio-Rad Laboratories (USA) | Cat#QX200 |
| PikoReal-Real time PCR system | Thermo Fisher Scientific (USA) | https://tools.thermofisher.com/content/sfs/manuals/PikoReal-Real-Time-PCR-System-User-Manual-v2.2.pdf |
| LAS-4000 mini imager | FUJIFILM | N/A |

## EXPERIMENTAL MODEL AND STUDY PARTICIPANT DETAILS

### Cell lines and primary airway epithelial cells

IB3-1 bronchial epithelial cells derived from a patient with cystic fibrosis carrying ΔF508/W1282X CFTR mutations were maintained as described previously.^14^ CF human primary bronchial epithelial cells, D-HBE-CF cells, were maintained as described previously.^42^ Cells were cultured at 37°C in a humidified atmosphere containing 5% CO_2_. Donor information for the D-HBE-CF cells is provided in the key resources table. CFTR mutation information for the D-HBE-CF cells was not provided by the supplier.

## METHOD DETAILS

### Natural-product library screening

An in-house natural-product library comprising 284 compounds derived from microorganisms and marine organisms was used for the primary screen. Microbial-derived compounds included secondary metabolites and SCFAs. Collection and use of microbial and marine materials were conducted in accordance with relevant guidelines and regulations, and the required approvals and collection permits were obtained. Most compounds were prepared as 2 mM stocks in dimethyl sulfoxide (DMSO) and stored at − 80°C, whereas SCFAs were dissolved in phosphate-buffered saline (PBS) before treatment. For screening assays, compounds were diluted immediately before use and applied for 24 h. Concentrations for SCFAs and non-SCFA compounds are indicated in the relevant figures and figure legends. The final DMSO concentration was kept constant across applicable conditions.

For hit selection, cutoff values were determined from the primary screening dataset of 284 compounds. The cutoff for WT-SIGIRR induction was defined as the mean relative WT-SIGIRR mRNA level plus two sample standard deviations, corresponding to mean + 2SD. The cutoff for WT-SIGIRR selectivity was similarly defined as the mean WT-SIGIRR/Δ8-SIGIRR mRNA ratio plus two sample standard deviations. Compounds exceeding these cutoffs were considered positive hits for WT-SIGIRR induction and selective WT-SIGIRR induction, respectively.

### Chemical treatments and cytokine stimulation

Cells were treated with the indicated reagents at the concentrations and for the durations specified in the figure legends. Reagents used for chemical treatments and cytokine stimulation are listed in the key resources table.

### RNA isolation, cDNA synthesis, and quantitative PCR

Total RNA was isolated from IB3-1 and D-HBE-CF cells, and cDNA was synthesized using the reagents listed in the key resources table. mRNA levels were quantified by real-time PCR using gene-specific primers listed in Table 1, as described previously.^14^ Expression values were normalized to 18S rRNA.

**Table 1.**
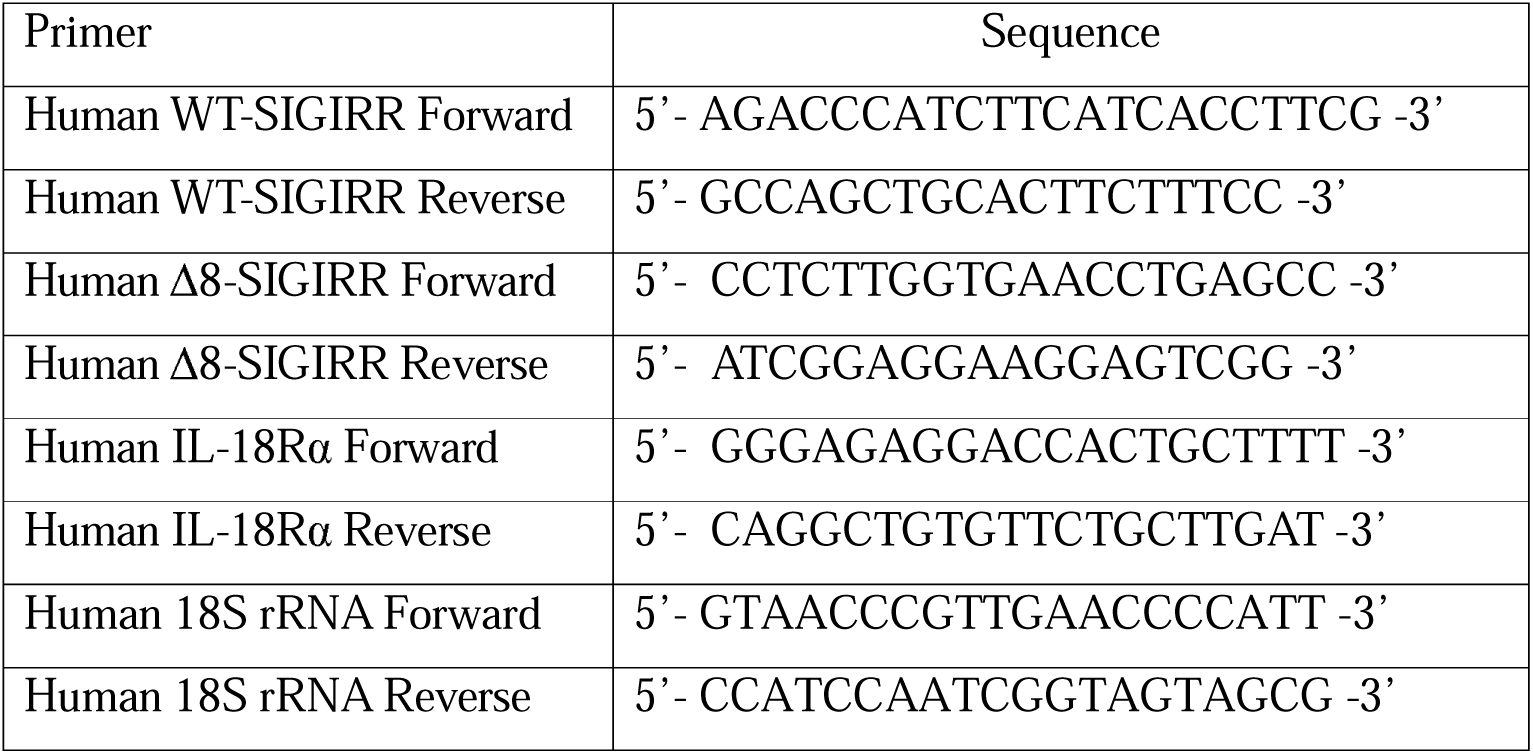
Sequences of primers for quantitative RT-PCR.

### siRNA-mediated HDAC3 knockdown

IB3-1 or D-HBE-CF cells were seeded in 12-well plates 24 h before transfection and cultured without antibiotics. Cells were transfected with 20 nM non-targeting control siRNA or HDAC3-targeting siRNA using the transfection reagent listed in the key resources table according to the manufacturer’s protocol. Cells were harvested 48 h after transfection unless otherwise indicated, and knockdown efficiency was confirmed by immunoblotting.

### Droplet digital PCR

WT-SIGIRR and Δ8-SIGIRR transcript copy numbers were measured by droplet digital PCR using the system and reagents listed in the key resources table according to the manufacturer’s protocol. Each reaction contained ddPCR supermix, custom assays for human WT-SIGIRR and Δ8-SIGIRR, cDNA, and nuclease-free water. Droplets were generated using an automated droplet generator, transferred to a 96-well PCR plate, and amplified under the following conditions: 95°C for 10 min; 40 cycles of 94°C for 30 s and 54°C for 1 min at a ramp rate of 2°C/s; and 98°C for 10 min. Droplets were read using a droplet reader and analyzed using the software listed in the key resources table.

### Expression constructs and transfection

Expression plasmids encoding C-terminally FLAG-tagged human WT-SIGIRR and IL-18Rα were generated in pCMV-FLAG-C and verified by Sanger sequencing. The expression plasmid encoding N-terminally HA-tagged human ubiquitin is listed in the key resources table. IB3-1 cells were seeded in 60-mm dishes at 5.0 × 10^5^ cells per dish and transfected at approximately 70%–80% confluence using the transfection reagent listed in the key resources table at a reagent-to-DNA ratio of 3:1. Cells were harvested 48 h after transfection unless otherwise indicated.

### Immunoblotting

Immunoblotting was performed as described previously.^43^ SIGIRR, IL-18Rα, HDAC3, LC3, β-actin, FLAG, and HA were detected using the antibodies listed in the key resources table. Signals were visualized using a chemiluminescent substrate and a luminescent image analyzer.

### Immunoprecipitation and ubiquitination assays

Cells were washed with PBS and lysed for 3 h at 4°C in RIPA buffer containing 1% NP-40, 0.1% sodium deoxycholate, 150 mM NaCl, and 50 mM Tris-HCl (pH 7.5), supplemented with protease inhibitors. Cleared lysates were incubated overnight at 4°C with control IgG or anti-FLAG antibody, followed by protein A/G agarose. Beads were washed five times with RIPA buffer, and bound proteins were eluted by boiling in SDS sample buffer.

For ubiquitination assays, cells expressing FLAG-tagged WT-SIGIRR or IL-18Rα together with HA-tagged ubiquitin were treated with RGFP966 as indicated. Immunoprecipitated proteins were analyzed by immunoblotting with anti-HA and anti-FLAG antibodies.

### Cell-surface biotinylation

Cells were washed with ice-cold PBS and incubated with a freshly prepared 1 mg/mL membrane-impermeable biotinylation reagent in cold PBS (pH 8.0) for 1 h on ice. Cells were then washed, lysed in RIPA buffer, and incubated overnight at 4°C with streptavidin agarose resin. Beads were washed six times, and biotinylated proteins were eluted with 4× SDS sample buffer.

## ELISA

Human IL-8 concentrations in culture supernatants were measured by ELISA using the kit listed in the key resources table, following the manufacturer’s protocol.

## QUANTIFICATION AND STATISTICAL ANALYSIS

Data are presented as mean ± SD unless otherwise indicated. Comparisons among multiple groups were performed by one-way analysis of variance followed by Tukey’s multiple-comparison test. *P* < 0.05 was considered statistically significant. The number of biological replicates is indicated in the figure legends. Statistical analyses and graph generation were performed using GraphPad Prism 11 listed in the key resources table.

