## Supplementary figures for "HDAC3 inhibition stabilizes the IL-37 receptor module to enhance anti-inflammatory signaling in cystic fibrosis airway epithelium"

### Figure S1

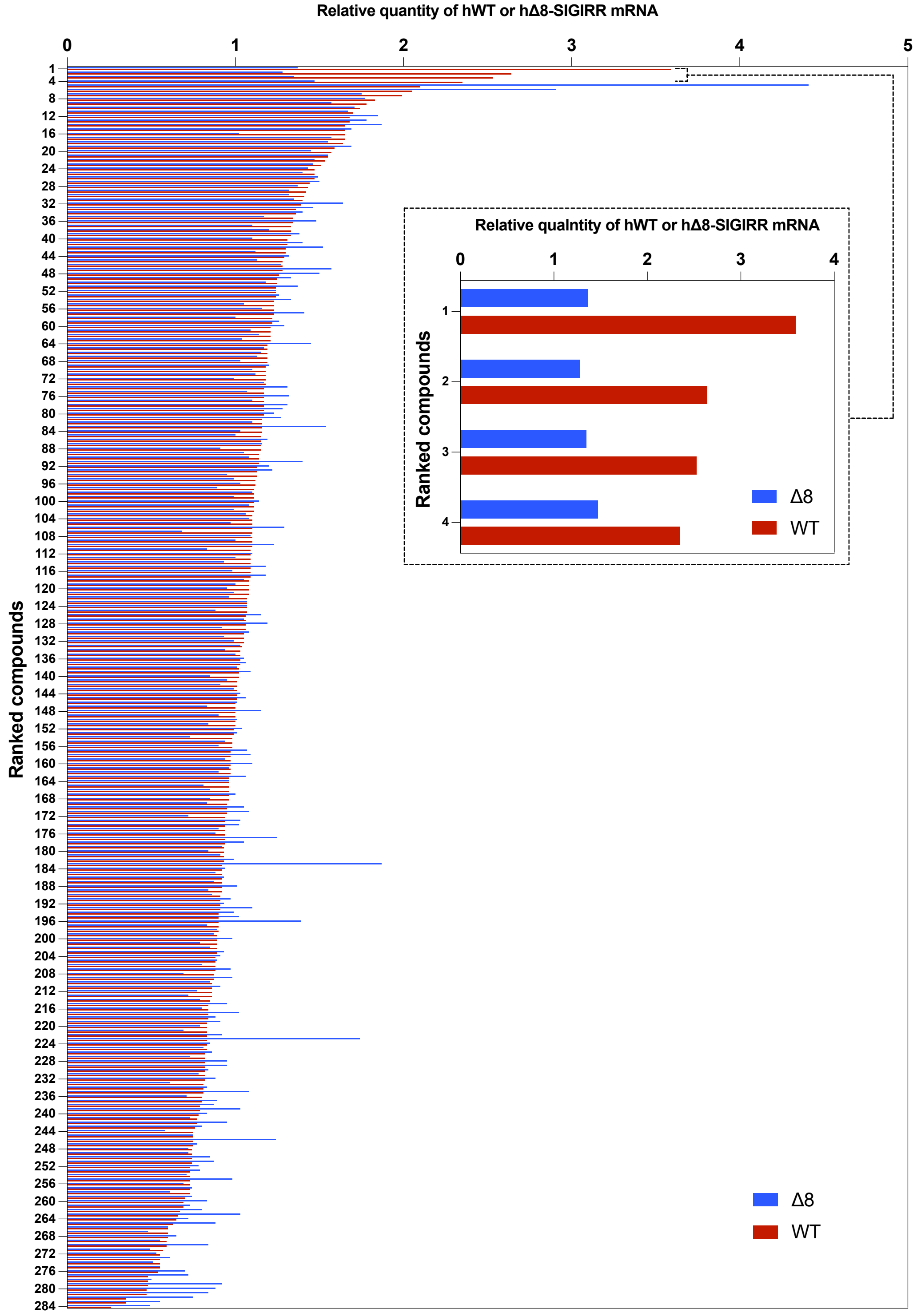


**Supplementary Figure S1. Screening of natural compounds that induce WT-SIGIRR and Δ8-SIGIRR mRNA expression in IB3-1 cells.**
IB3-1 cells were treated with 284 natural compounds at 10 μM for 24 h, and the relative mRNA expression levels of WT-SIGIRR and Δ8-SIGIRR were evaluated by quantitative RT-PCR. Compounds are ranked in descending order according to the induction level of WT-SIGIRR mRNA. Red bars indicate WT-SIGIRR mRNA expression, and blue bars indicate Δ8-SIGIRR mRNA expression. Values are shown as relative quantities normalized to the control condition. The inset shows an enlarged view of the top four WT-SIGIRR–inducing compounds. The top-ranked compounds were all short-chain fatty acids: 1, VA, valeric acid; 2, BA, butyric acid; 3, PA, propionic acid; and 4, AA, acetic acid.

### Figure S2

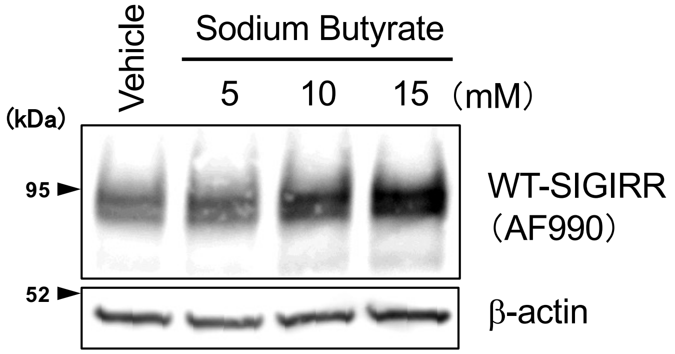


**Supplementary Figure S2. Sodium butyrate increases WT-SIGIRR protein abundance.**
IB3-1 cells were treated with the indicated concentrations of sodium butyrate, the sodium salt of butyric acid, for 24 h. WT-SIGIRR protein levels were analyzed by immunoblotting using an anti-SIGIRR antibody. β-actin was used as a loading control. Sodium butyrate increased WT-SIGIRR protein abundance, suggesting that BA-induced receptor upregulation is not simply attributable to nonspecific acidification. Representative blots are shown.

### Figure S3

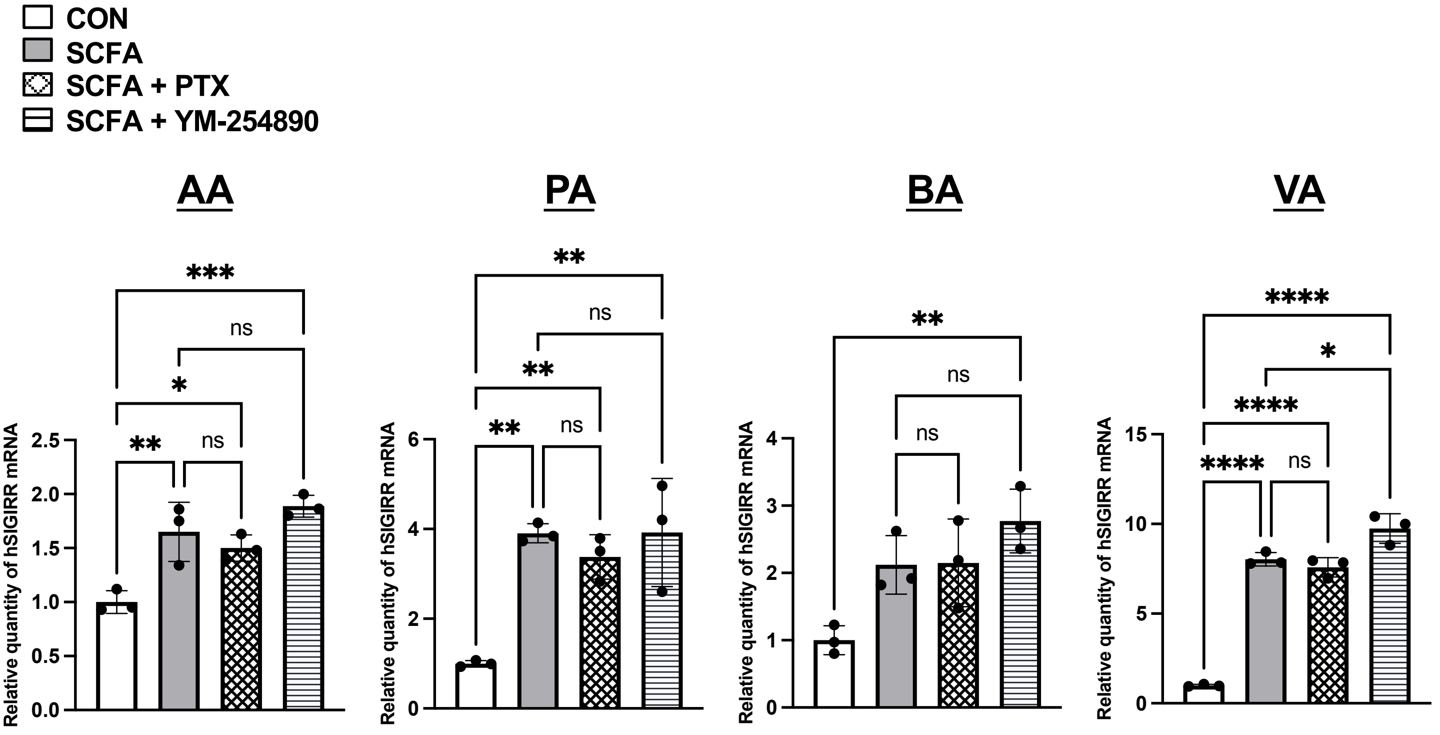


**Supplementary Figure S3. G protein-coupled receptor signaling inhibitors do not attenuate SCFA-induced WT-SIGIRR mRNA upregulation.**
IB3-1 cells were pretreated for 1 h with pertussis toxin (PTX), an inhibitor of Gi/o-coupled receptor signaling, or YM-254890, an inhibitor of Gq/11-mediated signaling. Cells were then treated with the indicated SCFAs—acetic acid (AA), propionic acid (PA), butyric acid (BA), or valeric acid (VA)—for 24 h in the continued presence or absence of each inhibitor. WT-SIGIRR mRNA expression was analyzed by quantitative RT-PCR and normalized to 18S rRNA. Data are presented as mean ± SD (n = 3). Statistical significance was assessed by one-way ANOVA followed by Tukey's multiple-comparison test. P values are indicated in the graphs; ns, not significant; *P < 0.05; **P < 0.01; ***P < 0.001; ****P < 0.0001. Neither PTX nor YM-254890 suppressed SCFA-induced WT-SIGIRR mRNA upregulation, suggesting that canonical Gi/o- or Gq/11-dependent signaling is dispensable for this response under the conditions tested.

### Figure S4

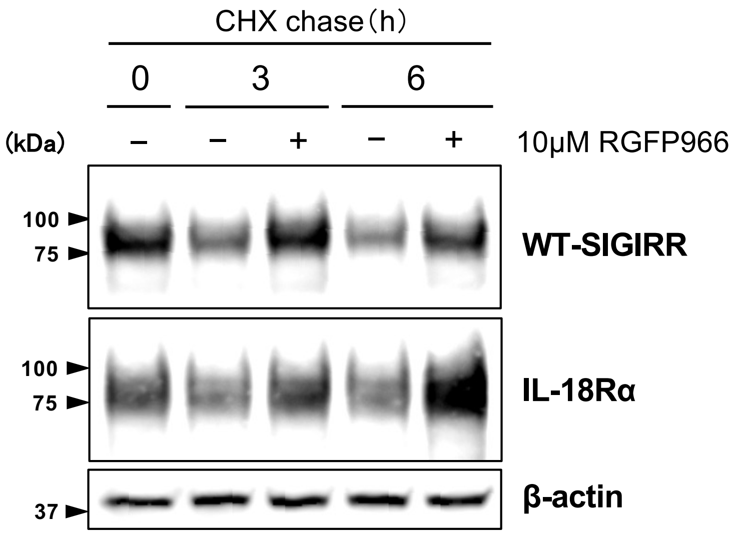


**Supplementary Figure S4. High-concentration RGFP966 stabilizes WT-SIGIRR and IL-18Rα during cycloheximide chase.**
IB3-1 cells were treated with cycloheximide (CHX; 100 μg/mL) to inhibit de novo protein synthesis in the presence or absence of RGFP966 (10 μM). Cells were harvested at the indicated time points, and WT-SIGIRR and IL-18Rα protein levels were analyzed by immunoblotting. β-actin was used as a loading control. RGFP966 delayed the decline in WT-SIGIRR and IL-18Rα protein abundance after CHX treatment, supporting a protein-stabilizing component of RGFP966-induced receptor upregulation. Representative blots are shown.

### Figure S5

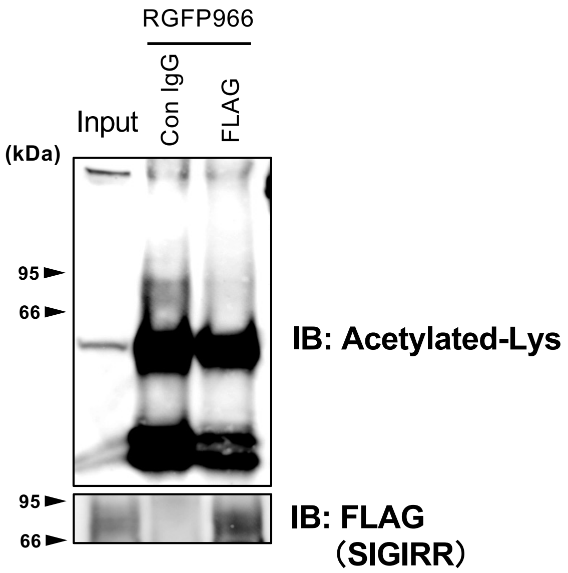


**Supplementary Figure S5. RGFP966 does not detectably increase lysine acetylation of WT-SIGIRR.**
IB3-1 cells expressing FLAG-tagged WT-SIGIRR were treated with RGFP966 (1 μM) for 24 h. Whole-cell lysates were subjected to immunoprecipitation with an anti-FLAG antibody, and lysine acetylation of immunoprecipitated WT-SIGIRR was assessed by immunoblotting with an anti-acetyl-lysine antibody. Immunoprecipitated WT-SIGIRR was confirmed by immunoblotting with an anti-FLAG antibody. Control IgG immunoprecipitation was included as a negative control. RGFP966 did not produce a detectable increase in lysine acetylation of WT-SIGIRR under these conditions. Representative blots are shown.
