## Supplementary material for "HDAC3 inhibition stabilizes the IL-37 receptor module to enhance anti-inflammatory signaling in cystic fibrosis airway epithelium": Source Data uncropped blots

### Original blot images

#### Figure 1D

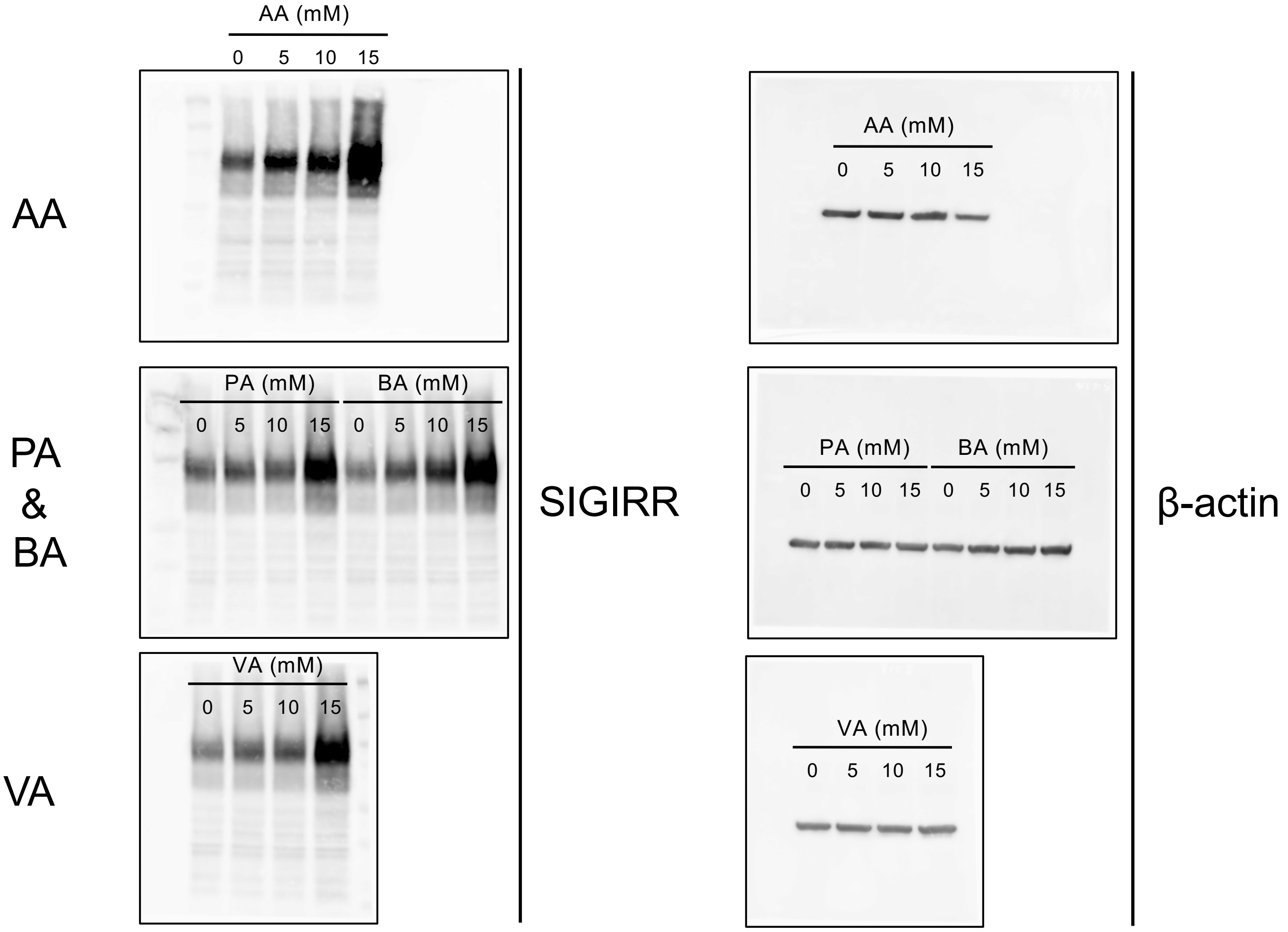


#### Figure 1 E

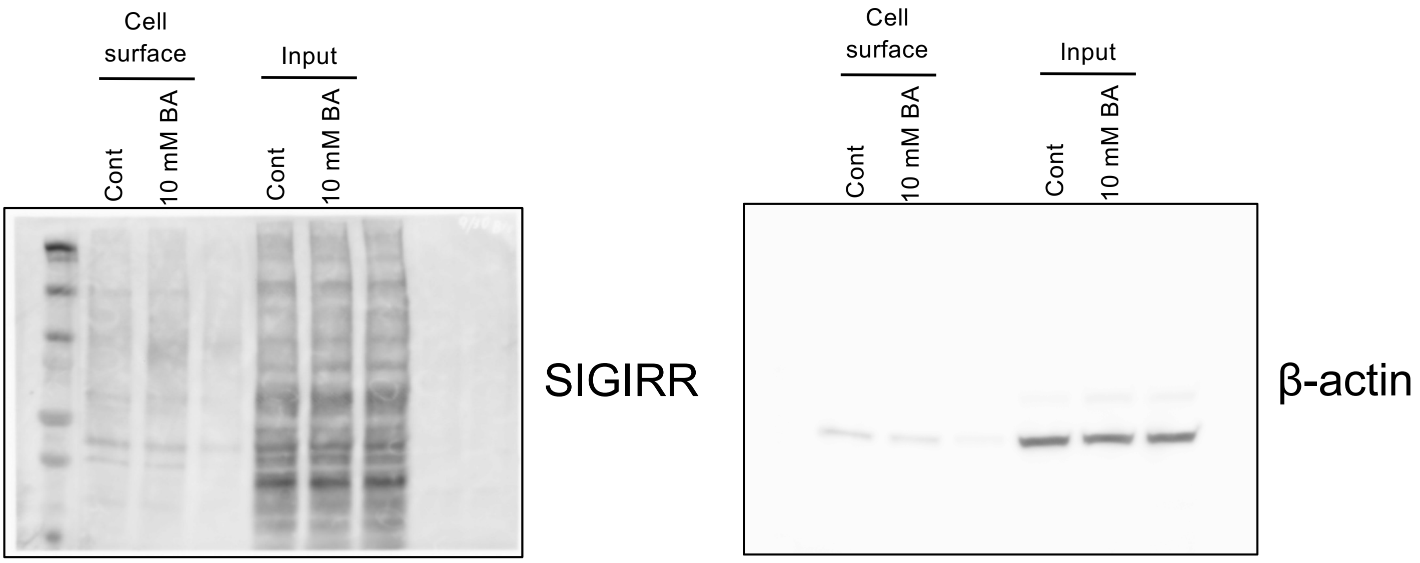

### Original blot images

#### Figure 2 B

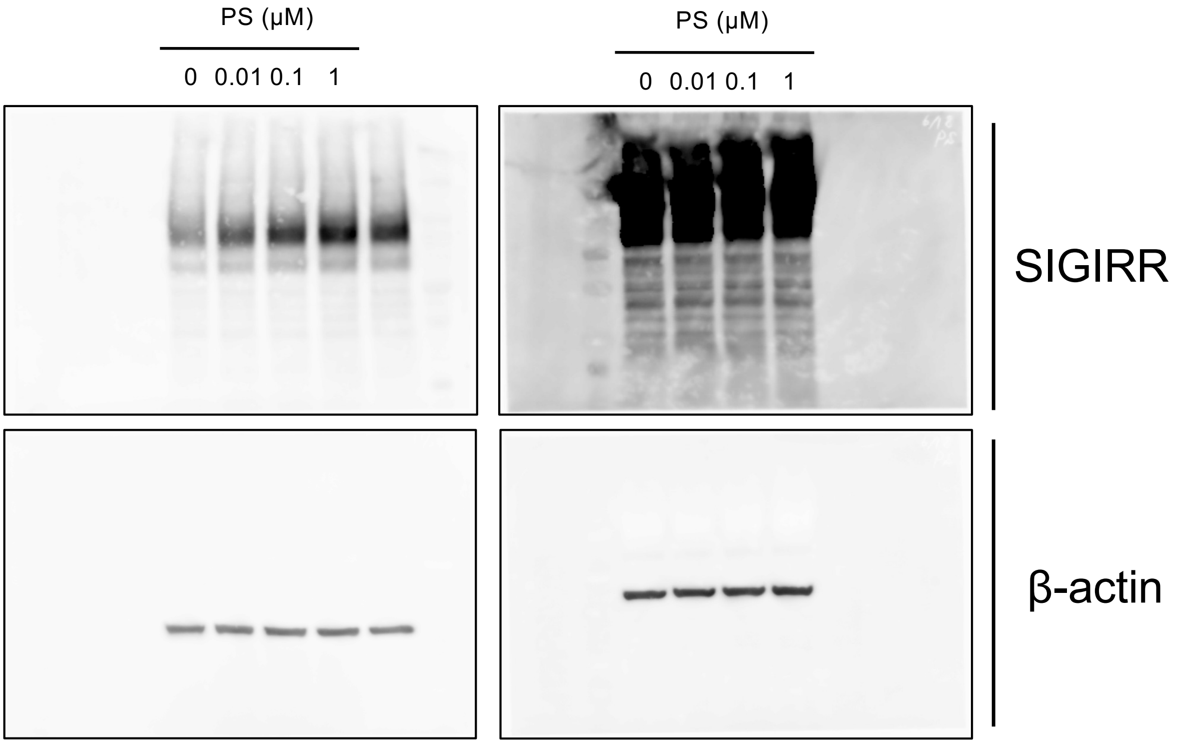

### Original blot images

#### Figure 4 A

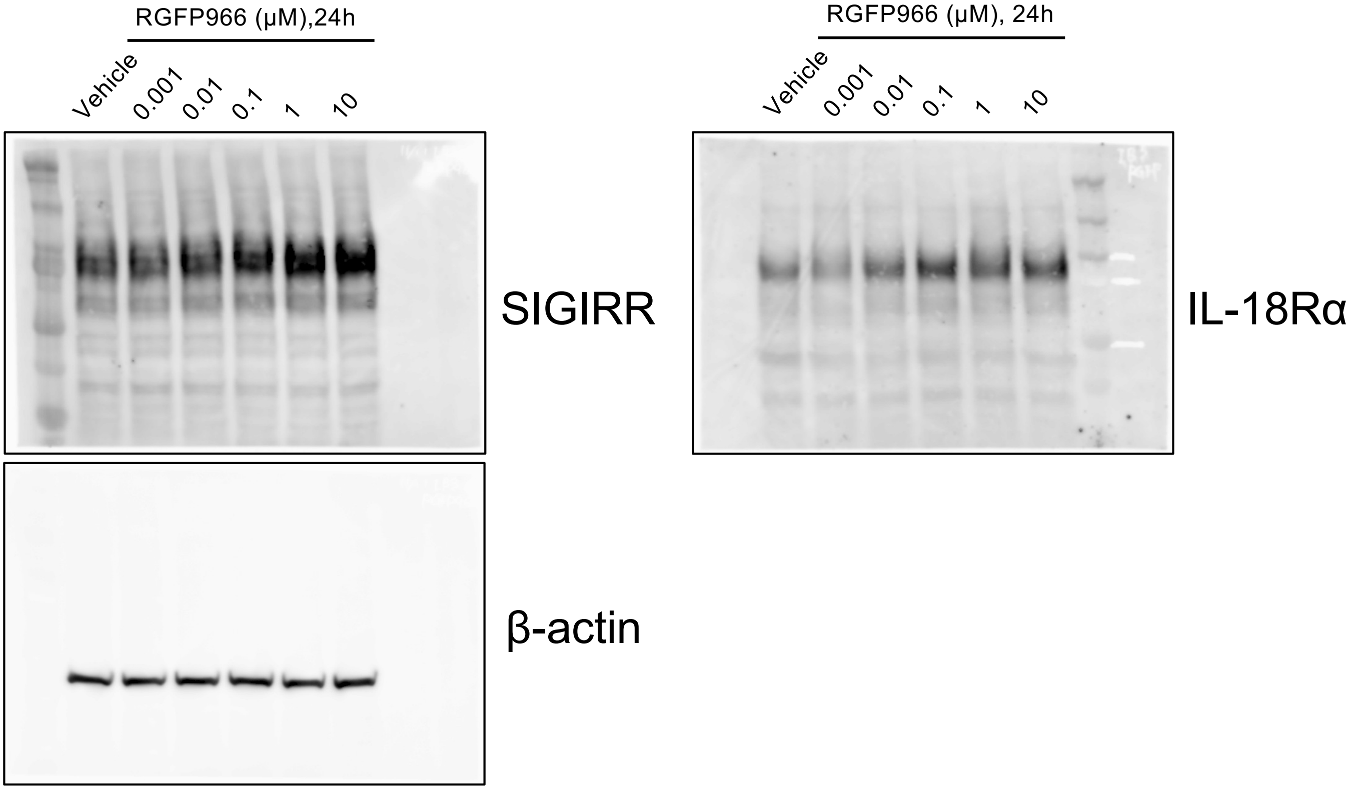


#### Figure 4 C

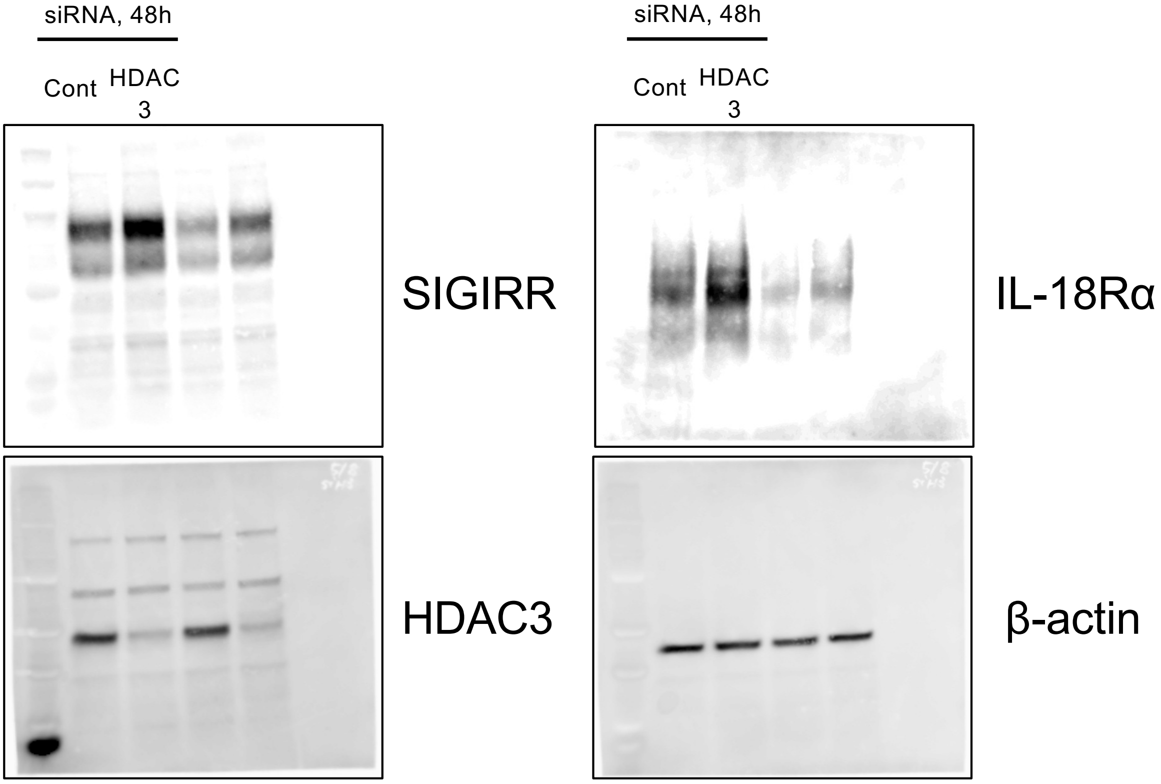

### Original blot images

#### Figure 4 E

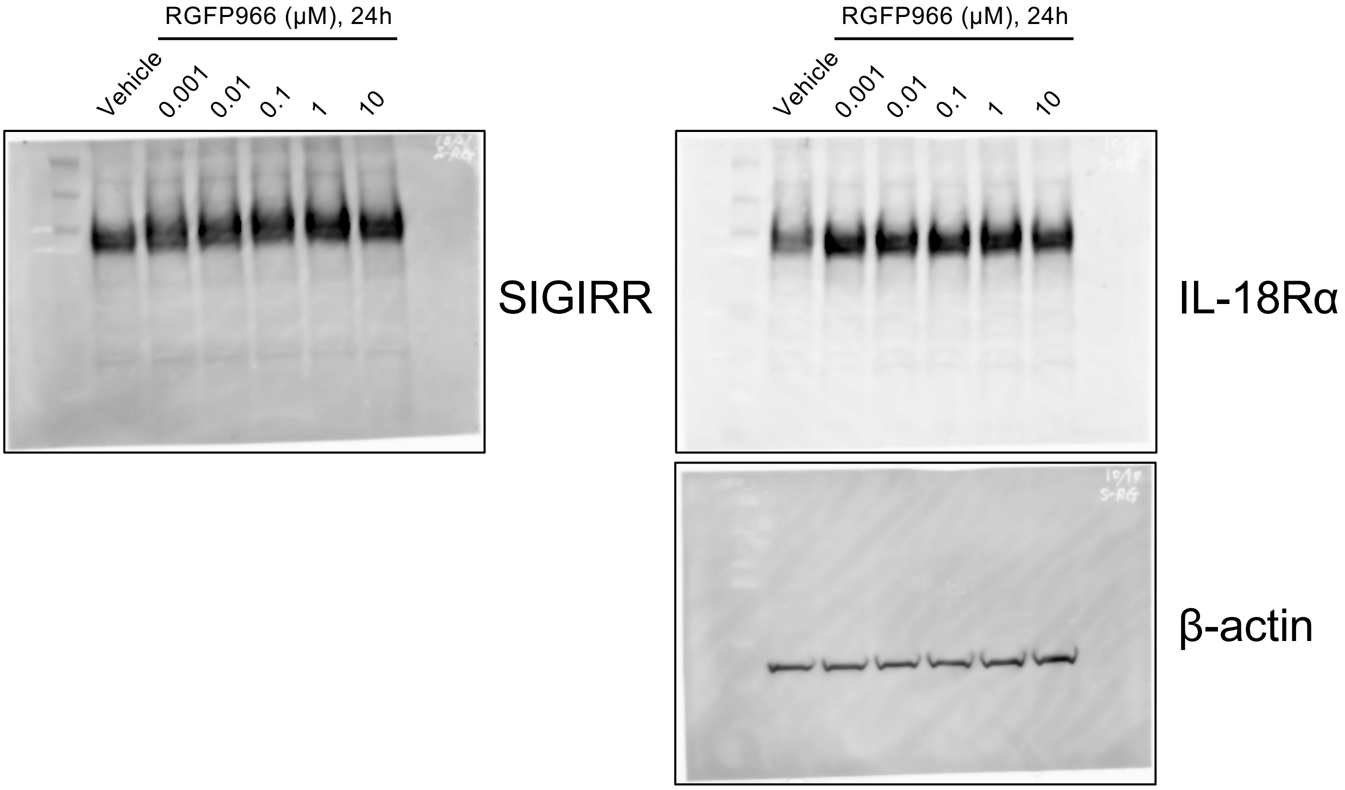


#### Figure 4 G

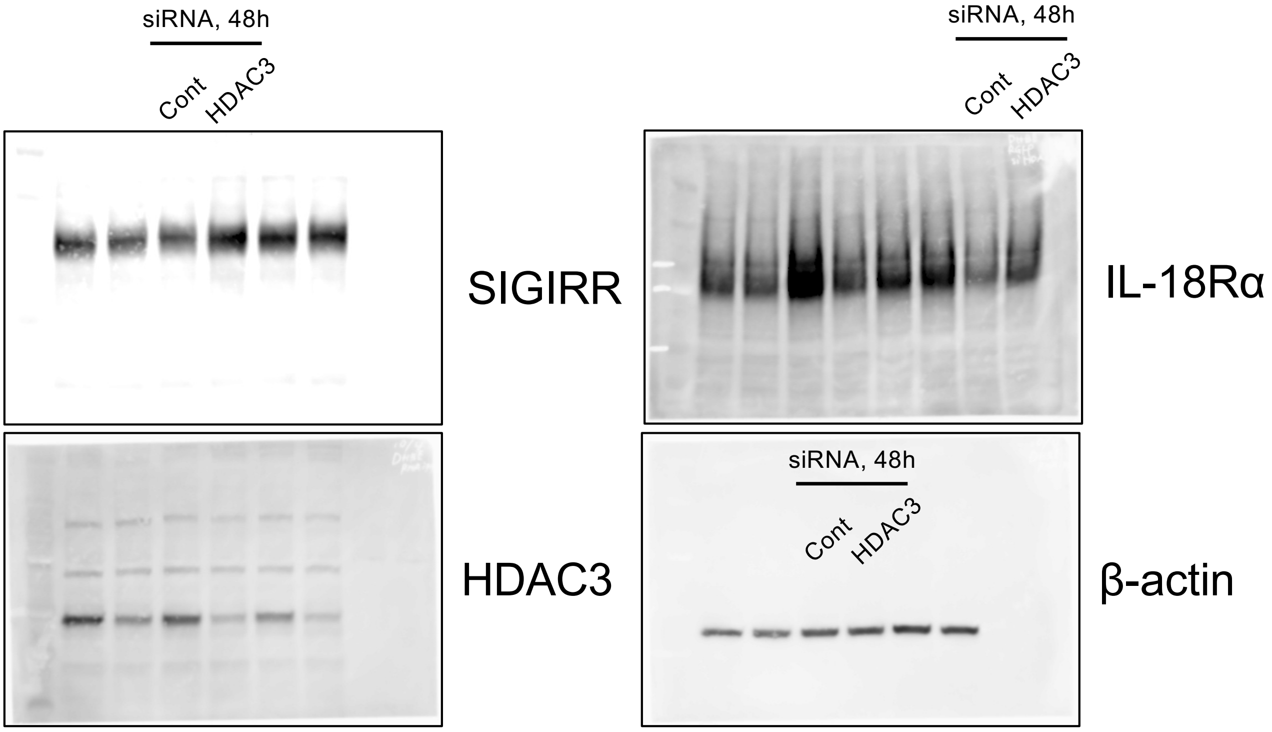

### Original blot images

#### Figure 5A

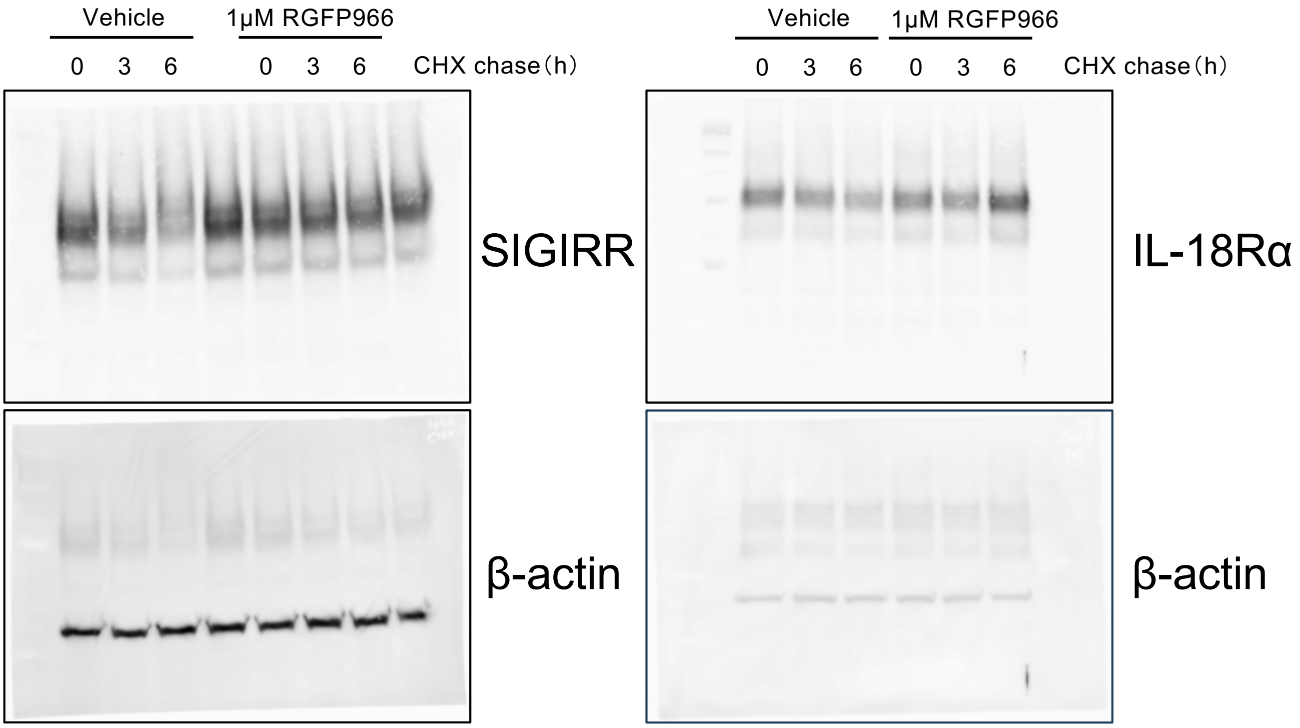


#### Figure 5B

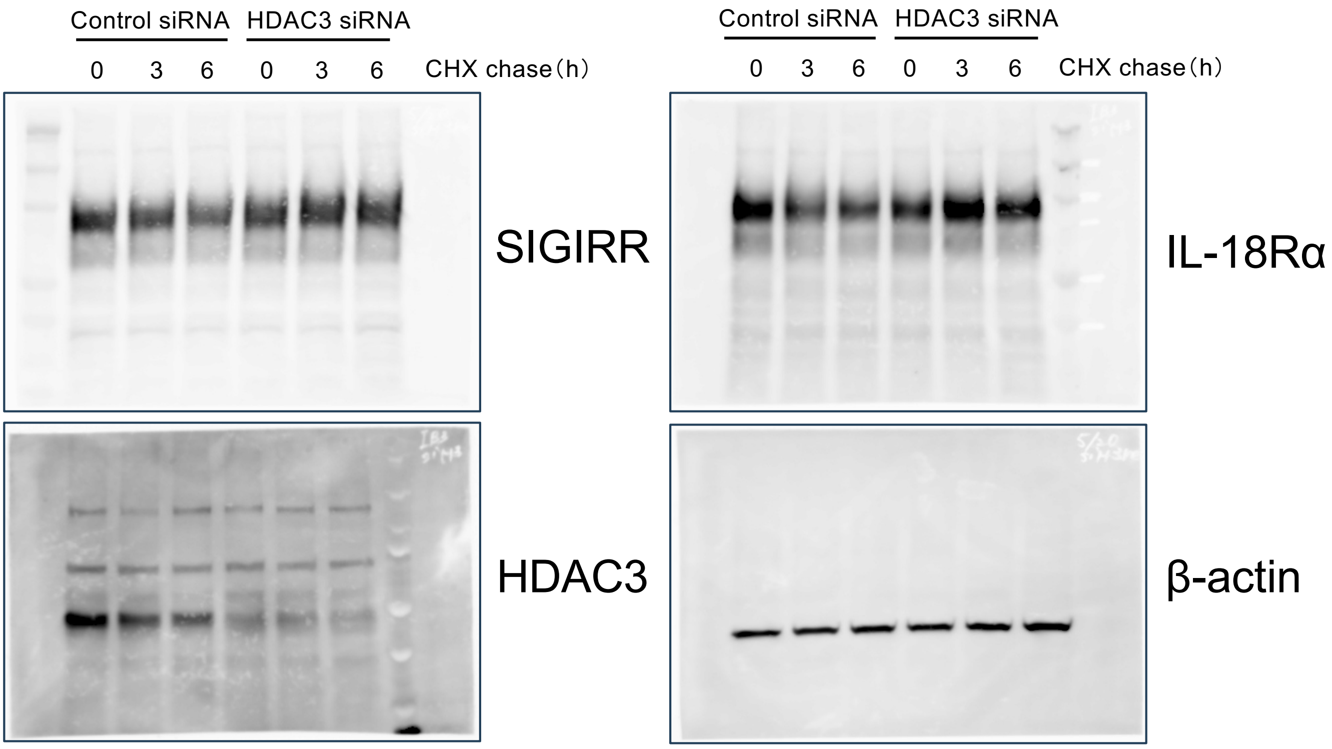

### Original blot images

#### Figure 5C

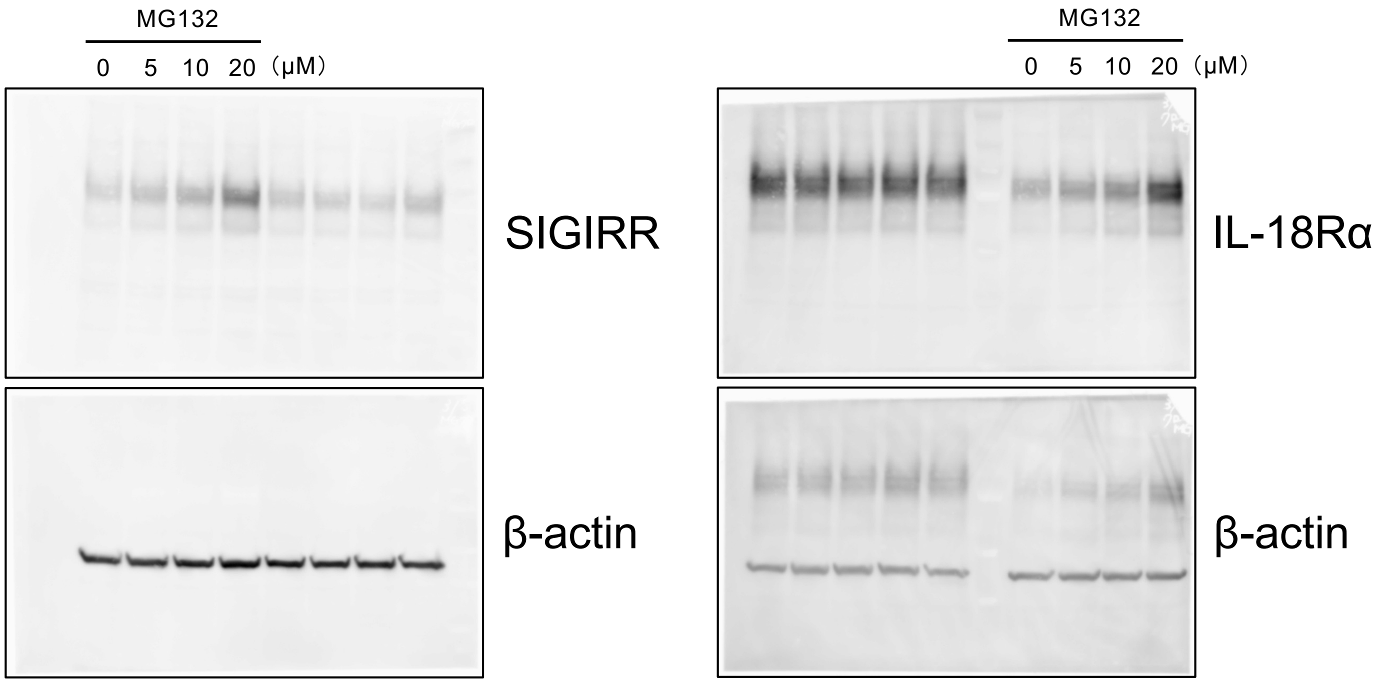


#### Figure 5D

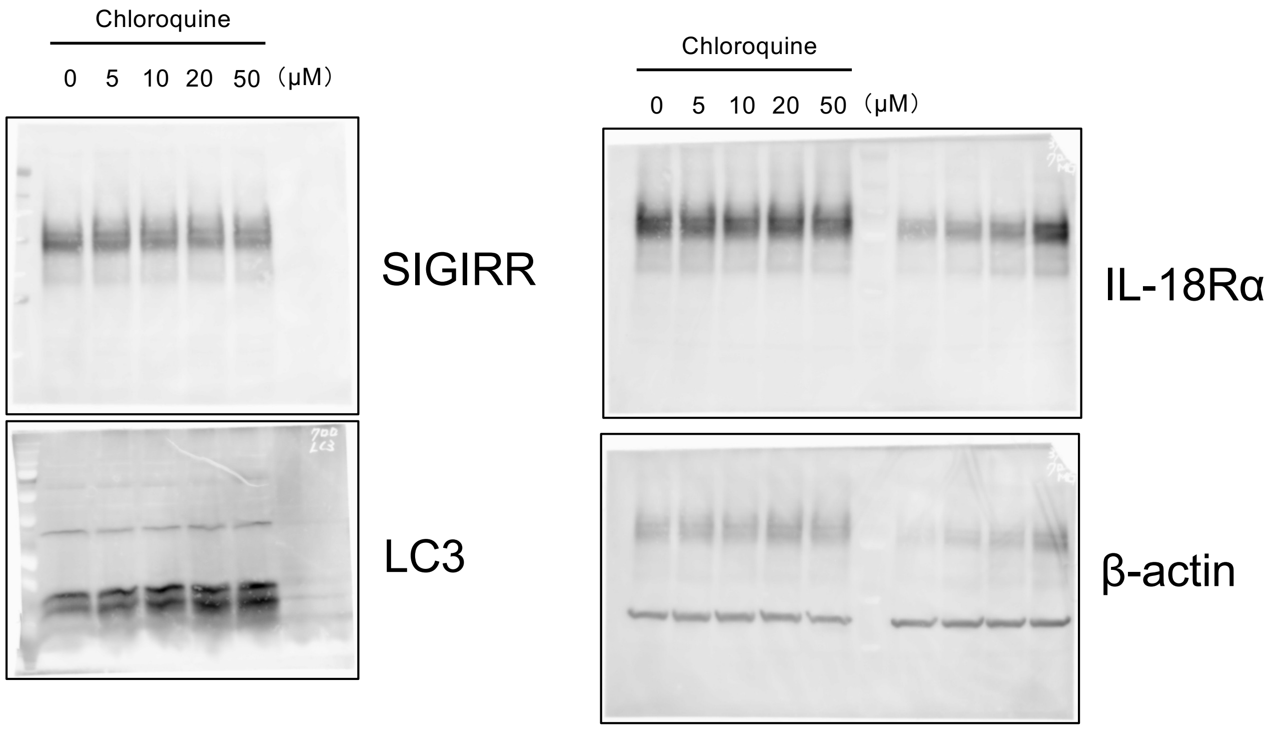

### Original blot images

#### Figure 5E

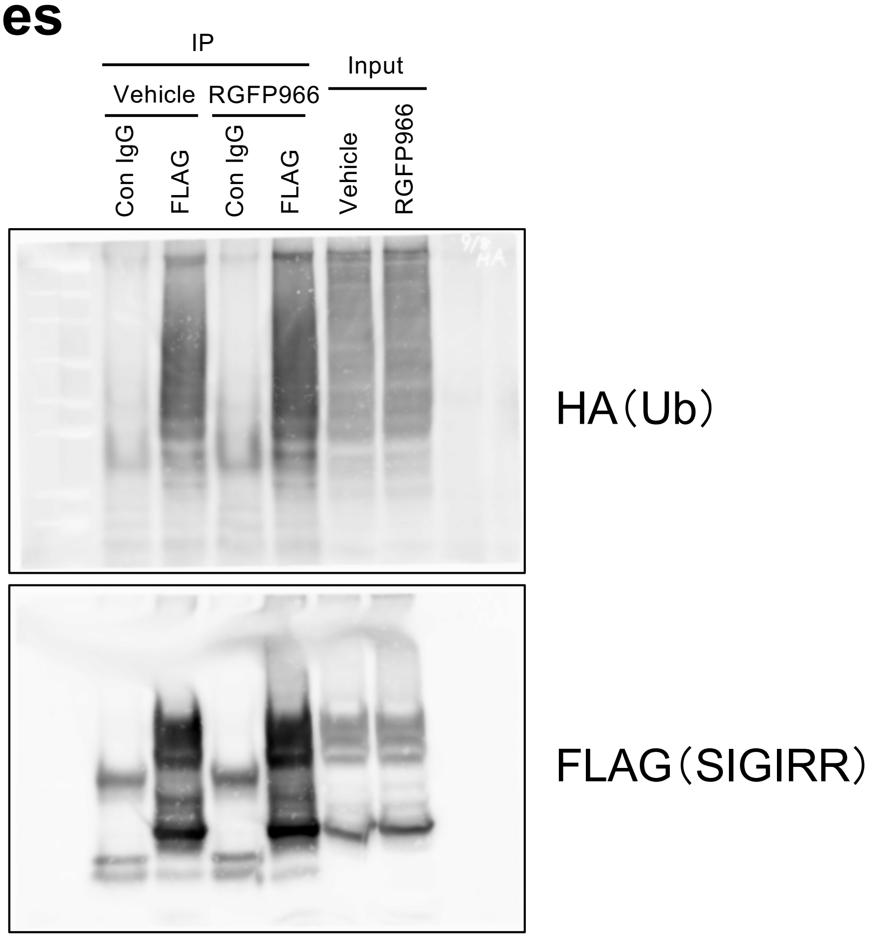


#### Figure 5F

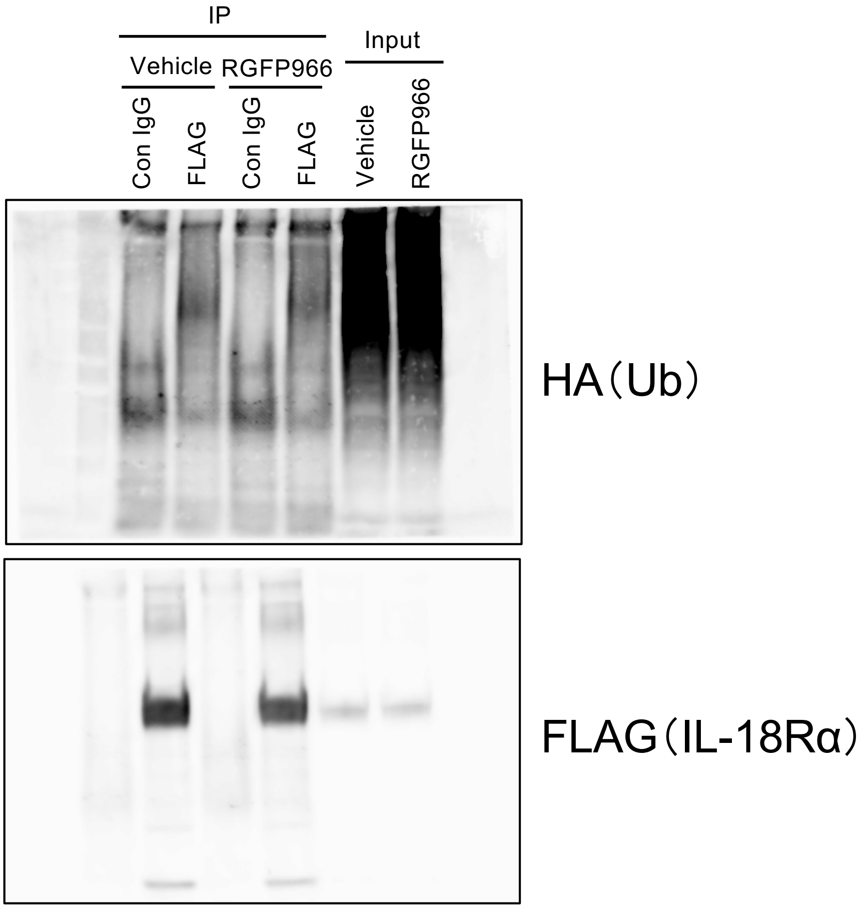

### Original blot images

#### Figure S2

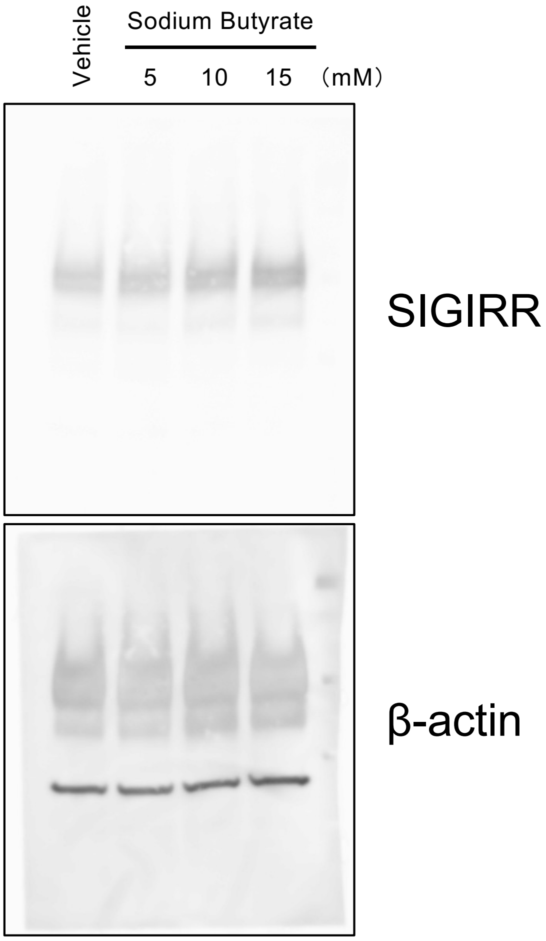


#### Figure S4

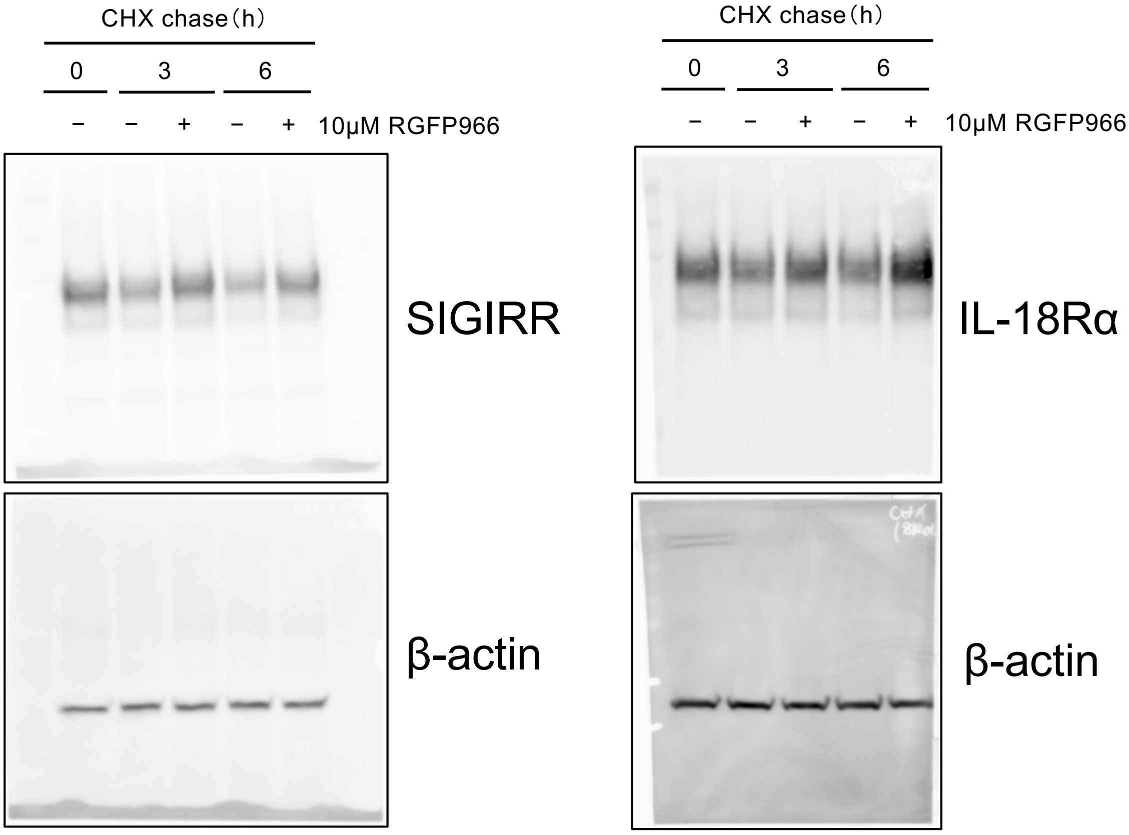
